# Genome-wide CRISPRi screens reveal the essentialome and determinants for susceptibility to dalbavancin in *Staphylococcus aureus*

**DOI:** 10.1101/2023.08.30.555613

**Authors:** Xue Liu, Vincent de Bakker, Maria Victoria Heggenhougen, Marita Torrissen Mårli, Anette Heidal Frøynes, Zhian Salehian, Davide Porcellato, Danae Morales Angeles, Jan-Willem Veening, Morten Kjos

## Abstract

Antibiotic resistance and tolerance remain a major problem for treatment of staphylococcal infections. Knowing genes that influence antibiotic susceptibility could open the door to novel antimicrobial strategies, including targets for new synergistic drug combinations. Here, we developed a genome-wide CRISPR interference library for *Staphylococcus aureus*, demonstrated its use by quantifying the essentialome in different strains through CRISPRi-seq, and used it to identify genes that modulate susceptibility to the lipoglycopeptide dalbavancin. By exposing the library to sublethal concentrations of dalbavancin using both CRISPRi-seq and direct selection methods, we found genes previously reported to be involved in antibiotic susceptibility, but also identified genes thus far unknown to affect antibiotic tolerance. Importantly, some of these genes could not have been detected by more conventional knock-out approaches because they are essential for growth, stressing the complementary value of CRISPRi-based methods. Notably, knockdown of a gene encoding the uncharacterized protein KapB specifically sensitizes the cells to dalbavancin, but not to other antibiotics of the same class, while knockdown of the Shikimate pathway surprisingly has the opposite effect. The results presented here demonstrate the potential of CRISPRi-seq screens to identify genes and pathways involved in antibiotic susceptibility and pave the way to explore alternative antimicrobial treatments through these insights.

## Introduction

The spread of antibiotic resistance in pathogenic bacteria combined with a halt in the antibiotic drug development pipeline represents a major threat to global health. Resistant *Staphylococcus aureus*, and particularly methicillin-resistant *S. aureus* (MRSA), is among the drug-resistant pathogens responsible for most deaths worldwide (1). For treatment of infections caused by resistant Gram-positive bacteria, glycopeptide antibiotics such as vancomycin or vancomycin-derivatives and the lipopeptide daptomycin are commonly used (2, 3). However, while being potent antimicrobial agents, resistance to these antibiotics is also rising (4). A continued effort to decipher mechanisms of resistance and identify novel drugs and combinatorial therapies as well as new drug targets is therefore critical. High-throughput genetics techniques linking genotypes to antibiotic susceptibility represent an attractive approach to identify potential novel combination therapies to combat resistance or resensitize antibiotic-resistant bacteria. For example, transposon insertion sequencing (Tn-seq) has been used to identify genetic determinants involved in susceptibility to daptomycin in *S. aureus* (5). One major limitation to such Tn-seq-based analyses, however, is that essential targets will not be included in the analyses as such mutants will not survive during library construction.

An approach to circumvent this is to perform genome-wide depletion of gene expression using CRISPR interference (CRISPRi). With CRISPRi, a catalytically inactive Cas9, known as dCas9, which can bind but not cleave DNA, is harnessed for repression of transcription (6, 7). dCas9 is co-expressed with a single guide RNA (sgRNA), a gene-specific RNA molecule that guides the dCas9 protein to its target sequence. The sgRNA consists of a 20-nt long target-specific sequence and a so-called Cas9 handle which is important for interaction with dCas9. By replacing the gene-specific sequence, the CRISPRi system can be directed to a new target. When bound within a gene, the dCas9-sgRNA complex acts as a transcriptional roadblock to block RNA polymerase, thereby inhibiting transcription (6, 7). Thus, CRISPRi will also inhibit expression of other genes co-transcribed with the target gene. CRISPRi has proven highly useful for functional studies of essential genes in a diversity of bacteria, including *S. aureus* (8–12). The relative simplicity and programmability of CRISPRi has also allowed upscaling to genome-wide CRISPRi libraries in bacterial species such as *Escherichia coli* (13–15), *Streptococcus pneumoniae* (16), *Streptococcus salivarius* (17), *Bacillus subtilis* (18), *Mycobacterium tuberculosis* (19), *Acinetobacter baumanii* (20) and *Vibrio natriegens* (21). Using a CRISPR adaptation strategy to generate guide RNAs, a CRISPRi library with an average of 100 sgRNAs targeting each gene, has also been developed for *S. aureus* (22). With such pooled CRISPRi libraries, the fitness of every single operon can be determined under different growth conditions by quantifying sgRNAs in the libraries with Illumina sequencing (termed CRISPRi-seq) (16, 23). CRISPRi-seq can be used to identify new targets for antibiotic therapy and we recently used CRISPRi-seq to show that amoxicillin-resistant *S. pneumoniae* can be resensitized to this antibiotic by combining it with the FDA-approved fertility drug clomiphene (24). In this work, we designed and developed a compact, genome-wide, inducible CRISPR interference library for *S. aureus* strain NCTC8325 and used this to identify essential genes and screen for genes influencing the susceptibility to the clinically relevant lipoglycopeptide antibiotic dalbavancin.

Dalbavancin is a semi-synthetic lipoglycopeptide with broad activity against Gram-positive bacteria including staphylococci, streptococci and enterococci (25). Like other glycopeptide antibiotics such as vancomycin and teicoplanin, dalbavancin targets cell wall synthesis by binding to the D-ala-D-ala dipeptide terminus of the lipid II, thereby sequestering this molecule, inhibiting peptidoglycan polymerization and crosslinking. Dalbavancin has a typical heptapeptide core which is modified compared to other glycopeptides and carries a lipid side chain. This side chain may be involved in its mechanism of action by anchoring the molecule to the bacterial cell membrane (26) and/or interacting with charged phospholipid head groups (27), although this has not been established. Dalbavancin is a highly potent anti-MRSA drug and has also been shown to be active against strains with reduced susceptibility to vancomycin or the last-resort lipopeptide drug daptomycin (28). The drug has been approved for use against acute bacterial skin and skin-structure infections (ABSSSIs) (29) but is also used for treatment of endocarditis and osteomyelitis (30). Dalbavancin has unique pharmacokinetic properties, with a half-life of more than a week, allowing for once-weekly administration of the drug (31, 32).

While still rare, dalbavancin non-susceptible mutants have been isolated from patients (33). One of these was shown to harbour mutations in *yvqF* (*vraT*), a membrane protein encoded in the same operon as the two-component system VraSR. VraSR is known to regulate cell wall metabolism (34) and has previously been associated with vancomycin-intermediate *S. aureus* (VISA) strains (35). *In vitro* exposure of *S. aureus* to dalbavancin has also been shown to select for tolerant mutants harbouring mutations in genes associated with VISA strains (35, 36). While vancomycin-resistant *S. aureus* (VRSA) strains are resistant to high glycopeptide concentrations due to the *vanA* resistance operon (2) causing direct target modification, VISA strains have a moderate reduction in susceptibility to glycopeptides. VISA is more prevalent than VRSA and represents a major problem for treatment of infections (35) and it is therefore critical to get better insights into the mechanism of action and understand which factors affect susceptibility in such cases. VISA strains mostly carry mutations that alter the cell surface to hinder the antibiotic from reaching its target and are often characterized by a thickened cell wall and altered transpeptidation cross-linking activity (35).

Here, we constructed a genome-wide CRISPRi library in the commonly used laboratory strain *S. aureus* NCTC8325-4, a prophage-rescued derivative of NCTC8325, and performed genome-wide CRISPRi-seq screens to quantify the essentialome and identify genes modulating the susceptibility to dalbavancin in *S. aureus*.

## Results

### Adapting the IPTG-inducible CRISPRi system in *Staphylococcus aureus* to facilitate library construction

In our previously described two-plasmid, IPTG-inducible CRISPRi system (12), *dcas9*-expression was controlled by a P_spac_ promoter containing a single *lacO* operator site in the plasmid pLOW-dcas9, while the sgRNA was constitutively expressed in plasmid pCG248-sgRNA(xxx) (12, 37). To reduce the background expression of *dcas9* in uninduced conditions, a second *lacO*-operator was inserted into the promoter to generate plasmid pLOW-P_spac2_-*dcas9* (**Fig. 1A**). To compare the dynamic range of induction for the two promoters, an *S. aureus* strain (MK1482) constitutively expressing GFP from the chromosome was used as a reporter. The *dcas9*-plasmids, as well as pCG248-sgRNA(*gfp*), expressing a *gfp*-targeting sgRNA, were transformed into the *S. aureus* GFP-reporter-strain and GFP-expression was measured with and without IPTG induction (90 min induction) (**Fig. 1B**). pLOW-P_spac2_-*dcas9* indeed showed a tighter regulation of *dcas9* compared to the strains carrying the original plasmid pLOW-*dcas9* in this reporter assay, and the former plasmid was therefore chosen for further use.

**Fig. 1.**
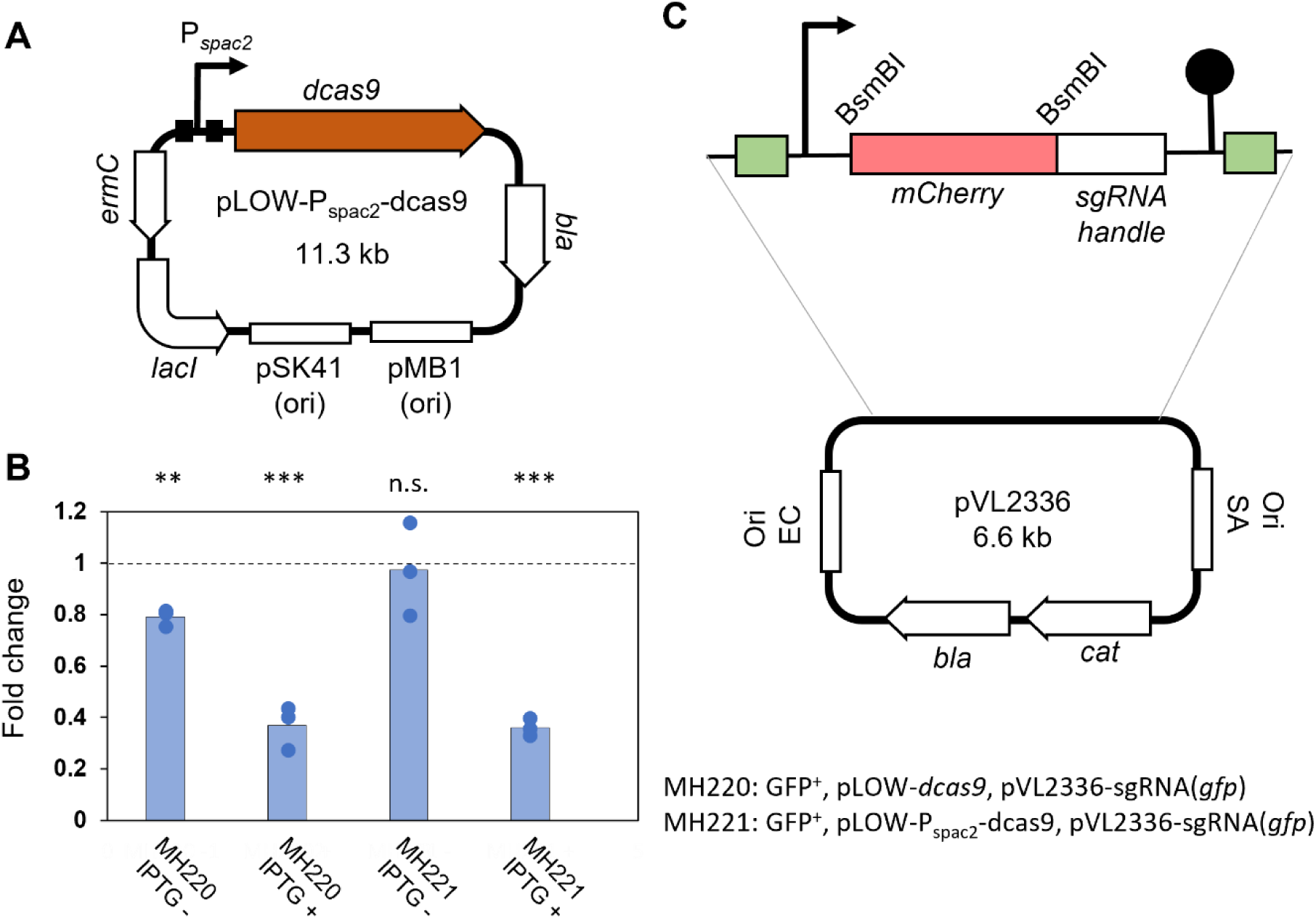
Adaptation of the staphylococcal CRISPRi system for CRISPRi-seq. **A**. Schematic representation of pLOW-P*_spac2_-dcas9*. The two *lacO* sites are indicated by black boxes. **B**. Reporter assay comparing background repression of GFP expression in CRISPRi-strains with the original system pLOW-dcas9 (in strain MH220) and the novel pLOW-P*_spac2_-dcas9* (in strain MH221). Exponentially growing cells were induced with 250 µM IPTG and relative fluorescence units (RFU)/OD were measured after 90 min depletion. Average fold-change in RFU/OD relative to a control strain constitutively expressing GFP (strain MK1482, indicated by dashed line) is plotted, along with data points from the three individual replicates. Two-sample t-tests, comparing each of the test strains to the control, was performed and significance is indicated by asterisks (p < 0.001), n.s.; not significantly different. **C**. Schematic representation of the pVL2336-plasmid. The green boxes represent the flanking Illumina amplicon sequences.

In our original CRISPRi system, we used inverse PCR to insert sgRNAs into the plasmid pCG248-sgRNA(xxx) (12). To aid the library construction, a new sgRNA vector (pVL2336) was engineered to allow insertion of sgRNAs by Golden Gate cloning using the class IIs restriction enzyme BsmBI (**Fig. 1C**). In pVL2336, two BsmBI restriction sites were inserted to flank an mCherry reporter gene placed exactly at the position of the 20-nt sgRNA base pairing region, thus allowing for red-white screening during Golden Gate cloning of new sgRNAs. In order to replace mCherry with 20-nt sgRNA base pairing regions, a forward and a reverse oligo, each of 24 nt (20-nt base pairing region and 4-nt overhangs to the digested plasmid), were designed and the two oligos were annealed and ligated into digested pVL2336, with a similar strategy as before (16). Furthermore, to enable one-step PCR during library preparation for amplicon sequencing, read 1, read 2, and adaptor sequences for Illumina sequencing were inserted in the sgRNA flanking sites.

### A genome-wide *S. aureus* CRISPRi library with 1928 sgRNAs provides wide coverage of multiple common strains

The genomic features and transcriptional units of *S. aureus* NCTC8325 defined by Mäder et al. (38) were used as a starting point for the design of sgRNA target sequences for a genome-wide *S. aureus* CRISPRi library. In total, 4028 genomic features have been annotated in the *S. aureus* NCTC8325 genome (accession number NC_007795/GCF_00013425), including ORFs, small RNAs and tRNAs. Genome-wide gene targeting in *S. aureus* is mainly restricted by the low transformation efficiency, which makes large sgRNA pools very challenging to construct. To get a concise, efficient, and balanced CRISPRi library, we decided to only include the ORFs and tRNAs in the library, resulting in 2836 features. The 1000 small RNAs were thus not directly covered in this study. Among the targeted features, 1041 are ORFs in transcription units (TU) encoded on the positive strand; 1159 features are ORFs in TUs on the negative strand and 636 features are not ascribed to any TU (38).

The following strategy was used for sgRNA design to cover all transcriptional units of the NCTC8325-4 genome (**Fig. 2A**): (i) An sgRNA was designed for all monocistronic ORFs; (ii) For the TUs with several ORFs, only the first ORF was selected as sgRNA target, since the same sgRNA will repress downstream genes due to polar effects of the CRISPRi approach; (iii) In addition, to reduce the risk of losing any genes in the library due to mis-annotation of the TUs, we introduced an extra sgRNA when the distance between two ORFs inside a predicted TU was more than 100 bp; (iv) For the 636 ORFs not ascribed to any TU, one sgRNA was designed for each of the individual ORFs (**Fig. 2A**).

**Fig. 2.**
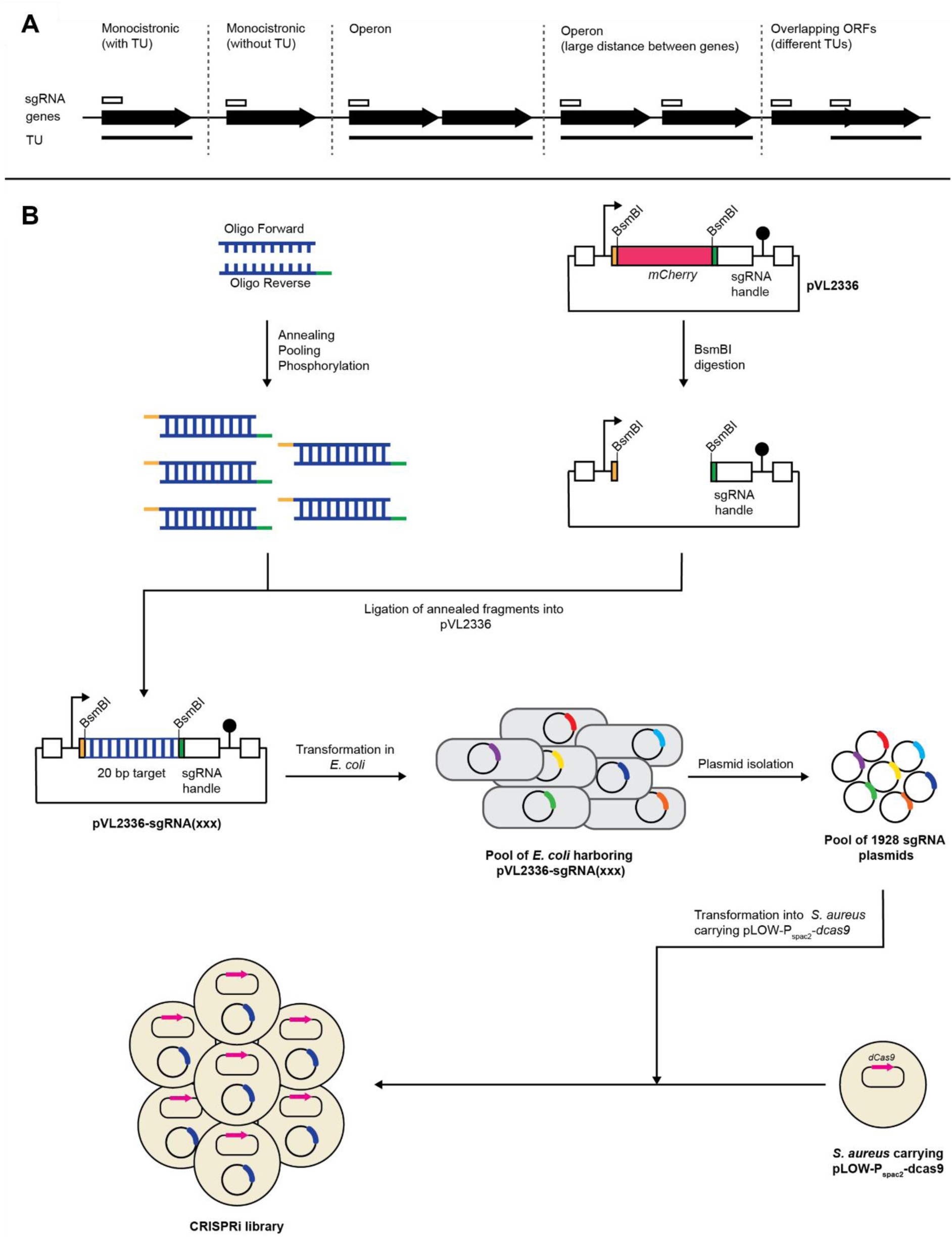
The *S. aureus* CRISPRi library. (A) Design strategy for sgRNAs in the CRISPRi library. Genes are indicated as arrows, transcriptional units (TUs) as black lines and sgRNAs as white boxes. (B) Schematic outline of the construction of the CRISPRi library. For each sgRNA, two 24-nt oligos were designed, with 4-nt overhang on each end to be compatible with the digested vector. The two oligos were annealed, and then the 1928 annealing products were pooled into one tube as the insert fragments for the following Golden Gate cloning. The plasmid pVL2336 was digested with BsmBI, and then the backbone was purified. The annealing pool was then ligated into digested pVL2336, followed by transformation into *E. coli* IM08B. The transformants were collected and pooled together as a reservoir for sgRNA plasmids. The sgRNA plasmids were then transformed into *S. aureus* with IPTG-inducible *dcas9*.

sgRNAs were designed to target the 5’ end of the selected genes (see Methods for details). In total, 1928 sgRNAs (**Table S1**) were designed to cover 2764 out of the total 2836 features (**Table S2**) in the NC_007795 genome used for sgRNA design (thus covering >97 % of all genomic features). The remaining 72 features in the NCTC8325 genome were not included in the library due to the lack of PAM sequences. Detailed information about the CRISPRi library, including a list of all sgRNA sequences (**Table S1**), an overview of all NCTC8325 locus tags along with corresponding sgRNA (**Table S2**) and a list of the non-targeted locus tags (**Table S3**) can be found in the supplemental information.

Our manually designed sgRNA library for NCTC8325 was evaluated for specificity and potential off-target effects according to de Bakker et al. (23). Eight sgRNAs were found to have more than one exact binding site in the genome, of which two not on the template strand within an annotated gene (**Table S1**). Specifically, sgRNA0034, sgRNA0179, sgRNA0929, sgRNA1456, sgRNA1900 and sgRNA1901 target conserved regions among gene sets SAOUHSC_00052-4, SAOUHSC_00269/00275/00276, SAOUHSC_01511/01512/01584, SAOUHSC_01410/01805/01881/02437, SAOUHSC_A01910/A01909 and SAOUHSC_A02013/00786, respectively. The former two sets are considered potential paralogs according to KEGG SSDB (39). The off-target analysis further revealed that some of these and a few other sgRNAs (sgRNA0268, sgRNA1294, sgRNA1580) have a relatively high expected off-target repression activity (**Table S1**), again partly due to potential paralogs. Some caution is therefore warranted when interpreting enrichment effects for these sgRNAs.

We also used the same pipeline to evaluate the sgRNA library functionality and genome coverage in seven other, commonly used *S. aureus* strains: Newman, JE2 USA300, COL, DSM20231, Mu50, RF122 and MRSA252 (**Table S4 – Table S10**). We estimate that the library covers 79-93% of all annotated features in those genomes, assuming similar operon structures and polar effects compared to the NCTC8325 strain the library was designed for (**Fig. S1A**). Note that this only concerns exact binding sites, meaning potentially even higher coverages when also considering those features with minor allelic differences, causing imperfect sgRNA binding in some strains. This indicates the library could potentially be used across different *S. aureus* strains.

To construct the CRISPRi library, we first made a plasmid pool with 1928 sgRNAs in *E. coli* IM08B. The 20-bp base-pairing region of the sgRNAs was first cloned into the above-mentioned *S. aureus* – *E. coli* shuttle vector, pVL2336, by pooled Golden Gate cloning, and then transformed into competent *E. coli* IM08B (**Fig. 2B**). In total, 1.3 × 10^6^ transformant colonies were obtained, providing a theoretical 674-fold coverage of the 1928-sgRNA library. The false-positive ratio of sgRNA cloning was estimated to be lower than 0.03% (no mCherry-positive colonies wcere identified among 3000 colonies). All the transformant colonies were pooled into one tube as a reservoir of sgRNA plasmids to construct the CRISPRi libraries in different *S. aureus* strains. To test the coverage and distribution of the 1928 sgRNAs, plasmids were purified from the pooled *E. coli* IM08B transformants and used as template in the PCR reaction for Illumina library preparation. Only one sgRNA was missing from the pool, confirming that the sgRNA plasmid pool was well-constructed, with sufficient coverage. Consistently, the distribution of the sgRNA counts confirmed that the library was well-balanced, as shown in **Fig. S2**. The plasmid pool was then transformed into *S. aureus* NCTC8325-4 carrying pLOW-P_spac2_-*dcas9* (**Fig. 2B**) in two independent experiments (Library 1 and 2). For each experiment, >200 000 colonies were collected and pooled for the final construction of the *S. aureus* CRISPRi library.

### CRISPRi-seq for genome-wide fitness quantifications in *S. aureus*

To test the potential of the constructed CRISPRi library, we first performed CRISPRi-seq to identify the essentialome in this strain. The library was grown in BHI medium in the presence or absence of IPTG for 12 (library 1) or 20 generations (library 2), plasmids were isolated and the sgRNA abundances were determined by Illumina sequencing (**Fig. 3A, Table S11**). Most of the CRISPRi strain heterogeneity between samples could indeed be attributed to *dcas9* induction (**Fig. S3**).

**Fig. 3.**
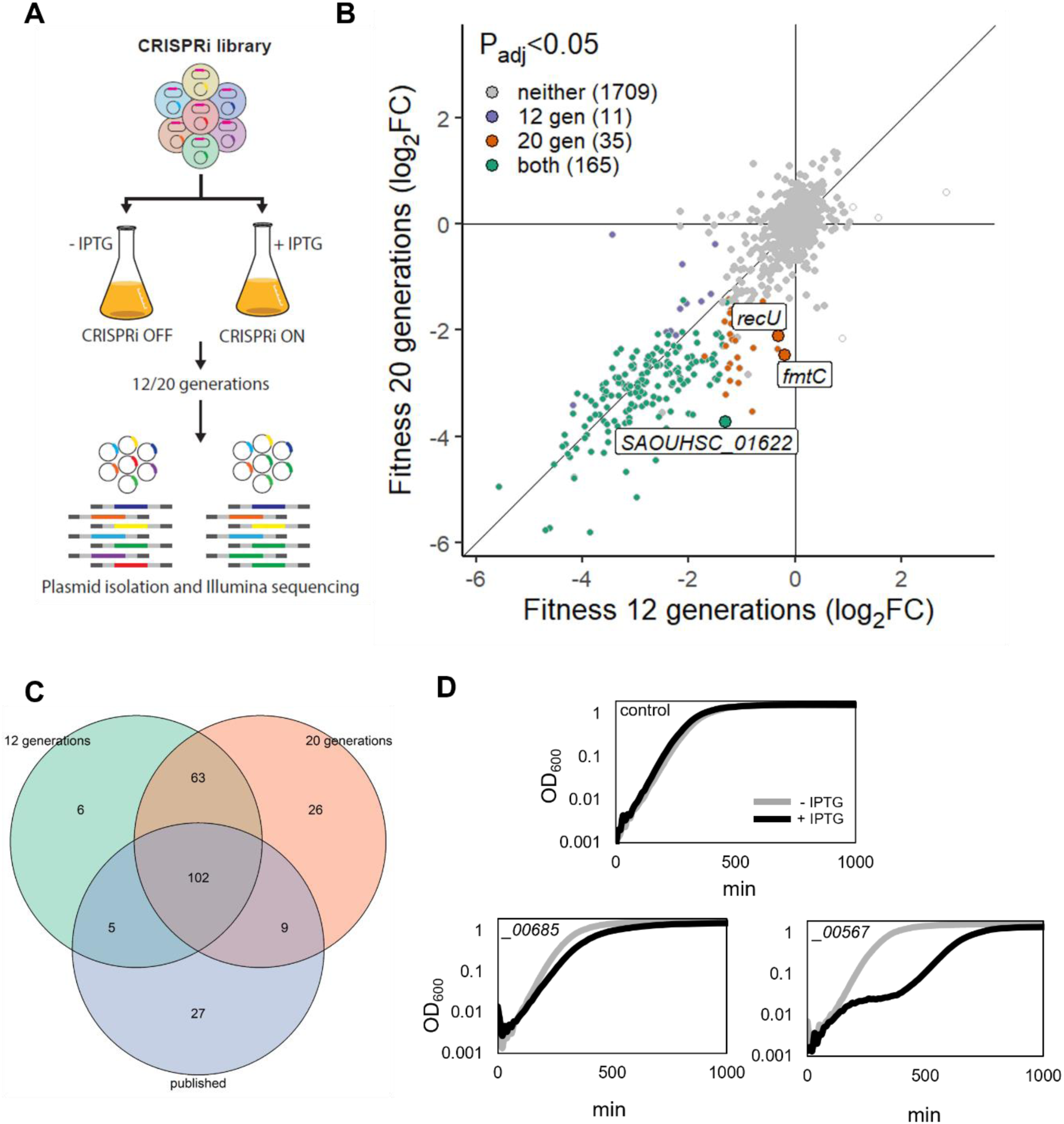
Essentialome of NCTC8325-4 as determined by CRISPRi-seq. (A) Flow-chart of the CRISPRi-seq screen. (B) Fitness effects upon *dcas9* induction for 12 versus 20 generations of exponential growth. Genes with significant differential fitness effects (|log_2_FC|>1, P_adj_<0.05) are depicted larger and alongside sgRNA target names. Points without color represent sgRNAs for which no p-value was determined due to extreme count outliers as detected by DESeq2. (C) Venn diagram showing comparison of essentialome with Tn-seq-based published essentialome data of related strains. (D) Experimental verification of newly identified essential genes in NCTC8325-4. ID numbers correspond to SAOUHSC_ locus tags. 500 µM IPTG was added for induction. The control strain harbours a non-targeting sgRNA (strain MM75).

We found 176 sgRNA targets to be essential after 12 generations of growth, and 200 after 20 generations (|log_2_FC|>1, P_adj_<0.05) (**Fig. 3B, Table S12**). Three sgRNAs were found to be significantly more depleted after 20 generations of growth, compared to 12 generations (|log_2_FC|>1, P_adj_<0.05) (targeting the genes *fmtC*, *recU-pbp2* and SAOUHSC_01622) (**Fig. 3B, Table S12**).

The *S. aureus* NCTC8325-4 essentialome, as we defined it by CRISPRi-seq, was compared with previous Tn-seq studies in other *S. aureus* NCTC8325-derived strains, including *S. aureus* HG001 grown in TSB (40) or BHI (41) and *S. aureus* SH1000 in BHI (42). Across these Tn-seq analyses, 212 genes were found to be essential across strains and growth media. These 212 conserved essential genes were targeted by 143 sgRNAs in our library, and we found that out of those, 75% (107 sgRNAs) and 78% (111 sgRNAs) were also determined to be essential by CRISPRi-seq in the 12- and 20-generation experiments respectively (**Fig. 3C**), indicating consistency between Tn-seq and CRISPRi-seq outcomes. Among the remaining sgRNAs expected to target essential genes, several have a relatively strong, but not significantly more than two-fold reduction (e.g., SAOUHSC_00009, SAOUHSC_00225 in the 20-generation experiment).

There are 63 sgRNA targets that are essential in both our CRISPRi-seq screens, but not conserved essential (**Fig. 3C**). However, 48 out of these are essential in at least one of the published essentiality screens (40–42). Thus, 15 sgRNAs in the CRISPRi-seq screen target genes which have not previously been defined as essential under rich growth conditions (**Table S13**). We suspected these might be explained by the well-documented polar effects of the CRISPRi system. As expected, nine sgRNAs are most likely defined as essential due to their effect on essential genes immediately up- or downstream of the target (**Table S13**). Of the six remaining sgRNAs, one targets *ccpA*, encoding the catabolite control protein CcpA which governs metabolic regulation, three target tRNAs, and two target genes of unknown function (SAOUHSC_00567 and SAOUHSC_00685). The essentiality of these last two genes was checked by individually constructed CRISPRi mutants, and knockdown of either gene resulted in a growth defect (**Fig. 3D**). We note that we cannot exclude that dCas9-related polar effects play a role in the essentiality of these newly identified putative essential genes, and this awaits future experimental validation.

Our sgRNA efficiency and specificity analysis showed that >89% of the sgRNAs are also functional in *S. aureus* Newman (**Fig. S1**). To show the potential of the sgRNA pool for multiple *S. aureus* strains, we performed a similar CRISPRi-seq screen in *S. aureus* Newman. A 12-generation CRISPRi-seq experiment was performed in the same way as shown in **Fig. 2A**. From this experiment, 202 sgRNA targets were defined as essential in *S. aureus* Newman (|log_2_FC|>1, P_adj_< 0.05) (**Fig. S4A, Table S14 - S15**). A total of 87% of the sgRNAs (153 sgRNAs out of 175) targeting an essential gene in NCTC8325-4 after 12 generations of growth were also found to have the same effect in Newman (**Fig. S4**). In contrast, CRISPRi-seq with the NCTC8325-4 library identified no sgRNA target as costly, whereas nine (|log_2_FC|>1, P_adj_<0.05) were in Newman, meaning that the depletion of these genes resulted in a relative increase in growth within the pooled library (**Fig. S4**). However, the costly phenotype of these genes was not confirmed with single knockdown strains; we tested three of these genes (*graX*, *vraF*, *saeR*) and the growth of the three CRISPRi knockdown strains did not show any significant difference from the control. Likely, the costly phenotype only reveals itself during competition and future experiments are required to test this hypothesis.

### CRISPRi-seq reveals factors important for susceptibility to the lipoglycopeptide dalbavancin

Dalbavancin is a lipoglycopeptide antibiotic whose primary mechanism of action is to block peptidoglycan polymerization via binding to the terminal D-ala-D-ala on the pentapeptide units (2). However, dalbavancin has some unique structural features compared to more thoroughly studied, related antibiotics such as teicoplanin, vancomycin (glycopeptides) and daptomycin (lipopeptide) (2, 43). We used CRISPRi-seq to get further insights into factors that affect susceptibility to dalbavancin (**Table S16 - S17**). The minimal inhibitory concentration (MIC) of dalbavancin against *S. aureus* NCTC8325-4 carrying the CRISPRi system (strain AHF10) was found to be 0.06 µg/ml. The induced CRISPRi library was then grown in the presence and absence of 0.015 µg/ml dalbavancin for 12 generations to identify sgRNAs that were over- or underrepresented after dalbavancin treatment (**Fig. 4A**). In total 27 genes were found to have a significant reduction in fold change (|log_2_FC|>1, P_adj_<0.05) in the presence of dalbavancin, indicating that knockdown of these genes results in increased sensitivity (**Table 1, Fig. 4B**), while 11 genes were found to have a significantly increased fold change under these conditions, suggesting that repression of these genes could cause reduced sensitivity to dalbavancin (**Table 1, Fig. 4B**).

**Fig. 4.**
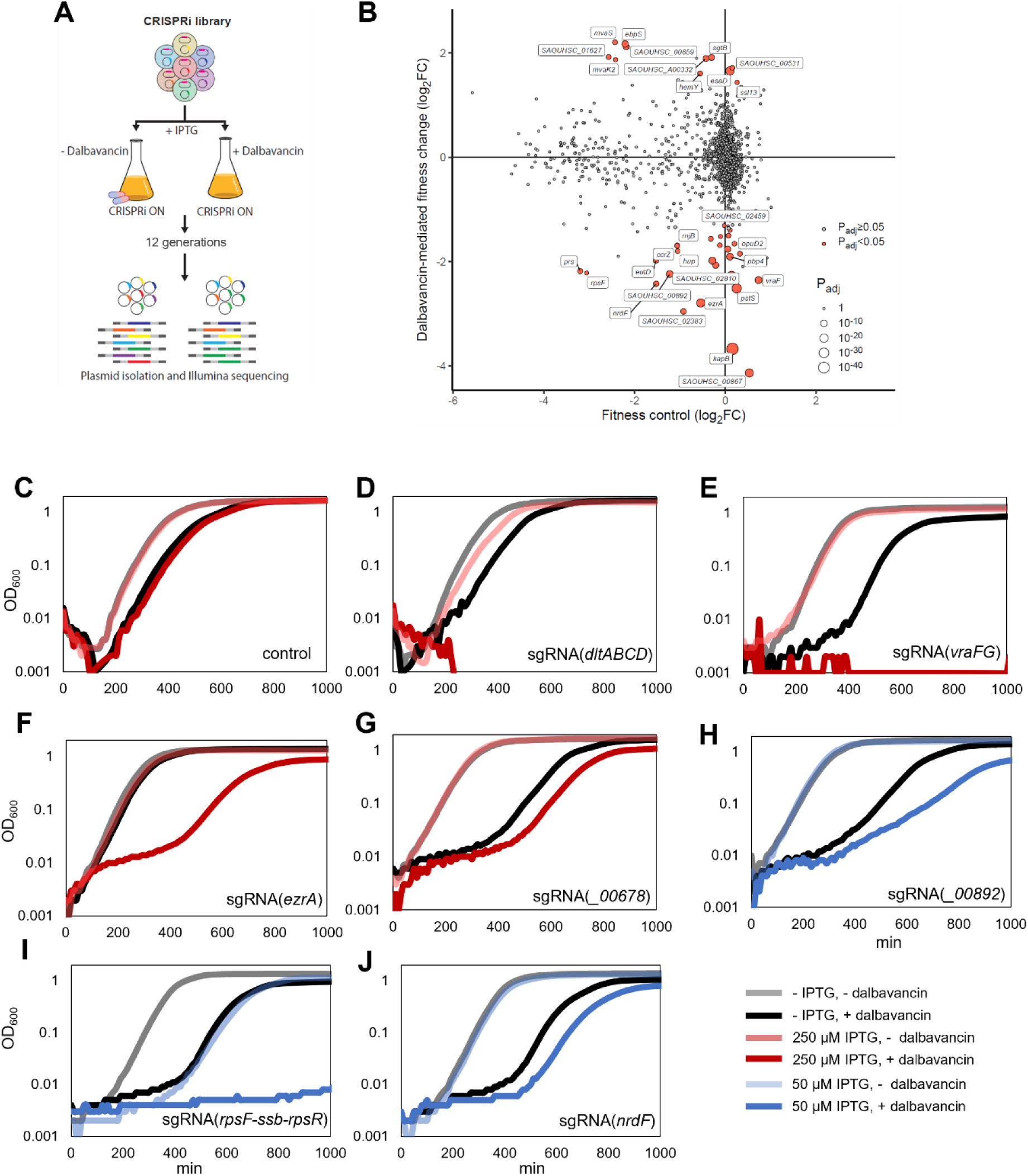
CRISPRi-seq identified factors affecting dalbavancin susceptibility. (A) Schematic workflow of the screen with dalbavancin stress. (B) CRISPRi-seq identified genes related to *S. aureus* sensitivity towards dalbavancin. Gene fitness was evaluated by CRISPRi-seq comparing CRISPRi-induced samples with a sublethal dose of dalbavancin to induced samples without dalbavancin (y-axis). This was contrasted with background fitness values as quantified with 12 generations of CRISPRi induction (x-axis, same as Fig. 3B). Targets of sgRNAs showing a significant change in fitness upon dalbavancin treatment were highlighted in red (|log_2_FC|>1, P_adj_<0.05). The top hits selected for confirmation were labelled with their targeted genes. (C-J) Confirmation of the screen with individual CRISPRi mutants. The CRISPRi system was induced with either 250 µM IPTG (red) or 50 µM IPTG (blue) to reach an appropriate degree of knockdown. Cells were grown with or without sublethal concentrations of dalbavancin (0.015 µg/ml for panels C, D and F or 0.03 for panels E, G, H, I and J). Non-targeting sgRNA-vector was used as control (C). Targeted genes were *dltA* (D), *vraFG* (E), *ezrA*, (F), SAOUHSC_00678 (G) SAOUHSC_ 00892 (H), *rpsF-ssb-rpsR* (I), *nrdF* (J).

**Table 1.** Genes affecting dalbavancin susceptibility as identified by CRISPRi-seq.

| CRISPRi target locus | Target genes | Function | log<sub>2</sub>FC | P<sub>adj</sub> | MIC |
| --- | --- | --- | --- | --- | --- |
| **SAOUHSC_00890** | ***kapB*** | **Unknown** | **-3.7** | **3.0E-47** | **0.045** |
| SAOUHSC_01389 | *pstS* | ABC transporter substrate-binding protein | -2.5 | 3.0E-27 | 0.06 |
| SAOUHSC_02810 | *SAOUHSC_02810* | Unknown | -2.3 | 5.4E-23 | 0.06 |
| **SAOUHSC_01827** | ***ezrA*** | **Septation ring formation regulator** | **-3.0** | **1.5E-20** | **0.03** |
| **SAOUHSC_00867** | ***_00867; dltXABCD*** | **D-alanylation of teichoic acids** | **-4.1** | **2.5E-19** | **0.03** |
| **SAOUHSC_02447** | ***_02447*** | **Unknown** | **-2.0** | **2.5E-12** | **0.045** |
| **SAOUHSC_00667** | ***vraFG*** | **ABC transporter** | **-2.4** | **4.7E-12** | **0.03** |
| **SAOUHSC_00892** | ***_00892*** | **Unknown** | **-2.2** | **6.2E-12** | **0.045\*** |
| **SAOUHSC_00646** | ***pbp4*** | **Penicillin binding protein 4** | **-1.9** | **7.1E-12** | **0.03** |
| **SAOUHSC_00678** | ***_00678*** | **Unknown** | **-2.1** | **4.3E-09** | **0.045** |
| SAOUHSC_01055 | *imp* | Inositol monophosphatase family protein | -1.8 | 1.2E-08 | n.t. |
| **SAOUHSC_02383** | ***_02383*** | **EVE-domain protein** | **-3.0** | **3.3E-06** | **0.045** |
| **SAOUHSC_00743** | ***nrdF*** | **ribonucleotide-diphosphate reductase** | **-2.4** | **5.9E-06** | **0.045\*** |
| SAOUHSC_00472 | *prs* | ribose-phosphate pyrophosphokinase | -2.2 | 3.3E-05 | 0.06 |
| SAOUHSC_00574 | *eutD; lipL* | Phosphate acetyltransferase_N-octanoyltransferase | -2.0 | 5.8E-05 | n.t. |
| SAOUHSC_01866 | *ccrZ* | cell cycle regulator | -1.7 | 0.00017 | n.t. |
| SAOUHSC_02444 | *opuD2* | BCCT-family osmoprotectant transporter | -1.8 | 0.00018 | n.t. |
| SAOUHSC_01466 | *recU; pbp2* | Holliday junction resolvase; penicillin-binding protein 2 | -1.6 | 0.00039 | n.t. |
| SAOUHSC_01548 | *_01548* | Conserved hypothetical protein | -1.5 | 0.00040 | n.t. |
| SAOUHSC_00833 | *_00833* | Nitroreductase domain protein | -1.7 | 0.00070 | n.t. |
| SAOUHSC_00663 | *_00663* | N-acetyltransferase domain-containing protein | -1.7 | 0.00070 | n.t. |
| SAOUHSC_01490 | *hup* | DNA-binding protein HU | -1.8 | 0.00071 | n.t. |
| SAOUHSC_00347 | *_00347* | Unknown | -1.4 | 0.00196 | n.t. |
| SAOUHSC_01252 | *rnjB* | Ribonuclease J2 | -1.7 | 0.00436 | n.t. |
| SAOUHSC_02740 | *_02740* | Drug transporter | -1.5 | 0.02159 | n.t. |
| **SAOUHSC_00348** | ***rpsF; ssb; rpsR*** | **Ribosomal; ssDNA-bind; ribosomal** | **-2.2** | **0.04321** | **0.045\*** |
| SAOUHSC_02459 | *_02459* | Unknown | -1.3 | 0.04388 | n.t. |
| SAOUHSC_02860 | *mvaS* | HMG-CoA synthase | 2.2 | 0.00016 | 0.06 |
| **SAOUHSC_01501** | ***ebpS*** | **cell surface elastin binding protein** | **2.2** | **2.52E-10** | **0.09\*** |
| **SAOUHSC_00659** | ***_00659*** | **Unknown** | **2.1** | **3.78E-08** | **0.09\*** |
| SAOUHSC_01627 | *_01627; _01628* | Unknown | 1.9 | 0.00015 | n.t. |
| SAOUHSC_02012 | *sgtB* | Monofunctional glycosyltransferase | 1.9 | 3.78E-08 | 0.06 |
| SAOUHSC_A00332 | *_A00332* | Unknown | 1.9 | 1.76E-07 | n.t. |
| SAOUHSC_00579 | *mvaK2* | phosphomevalonate kinase | 1.9 | 0.00978 | 0.06 |
| SAOUHSC_00531 | *_00531* | Unknown | 1.7 | 0.00040 | n.t. |
| SAOUHSC_00685 | *_00685; _00686* | Transcription factor; unknown | 1.7 | 0.12676 | 0.06 |
| SAOUHSC_00268 | *esaD* | Type VII secretion system EssD | 1.7 | 1.66E-20 | n.t. |
| SAOUHSC_01960 | *hemY* | protoporphyrinogen oxidase | 1.6 | 0.00026 | n.t. |

MIC-values are given in µg/ml. The MIC of the CRISPRi no-target control was 0.06 µg/ml. Experimentally verified hits are highlighted in bold. n.t.; not tested. \* essential genes affecting dalbavancin susceptibility.

We created a selection of 13 individual CRISPRi strains expected to have an increased sensitivity to dalbavancin and tested their susceptibility to dalbavancin in two-fold dilution assays in microtiter plates. Indeed, 10 out of 13 strains tested displayed increased susceptibility to dalbavancin in these assays, with a maximum two-fold MIC reduction (**Table 1**). These include genes or operons of known function, such as *dltABCD, vraFG*, *ezrA*, *rpsF*-operon, *nrdF* and *pbp4* (**Fig. 4C, D, E, F, I, J, Fig. 5A**), but also yet uncharacterized genes (e.g., SAOUHSC_00678, SAOUHSC_00892, *kapB*, **Fig. 4G, H** and **Fig. 5E**). Importantly, the verified hits identified in this screen include essential genes, which could not have been identified with any gene knockout approach due to their essentiality (**Fig. 4B**). For example, downregulation of *nrdF*, encoding a ribonucleotide-diphosphate reductase, SAOUHSC_00892, a putative RNA-binding protein of unknown function, as well as the ribosomal operon *rpsF-ssb-rpsR*, led to increased sensitivity to dalbavancin (**Fig. 5C-J**). To further corroborate the results of the screen, we also created deletion mutants of the two non-essential genes *kapB* and *pbp4*. Indeed, similar to the knockdowns (**Fig. 5A and E**), the deletion mutants (**Fig. 5B and F**) were hypersusceptible to dalbavancin, with a two-fold reduced MIC.

**Fig. 5.**
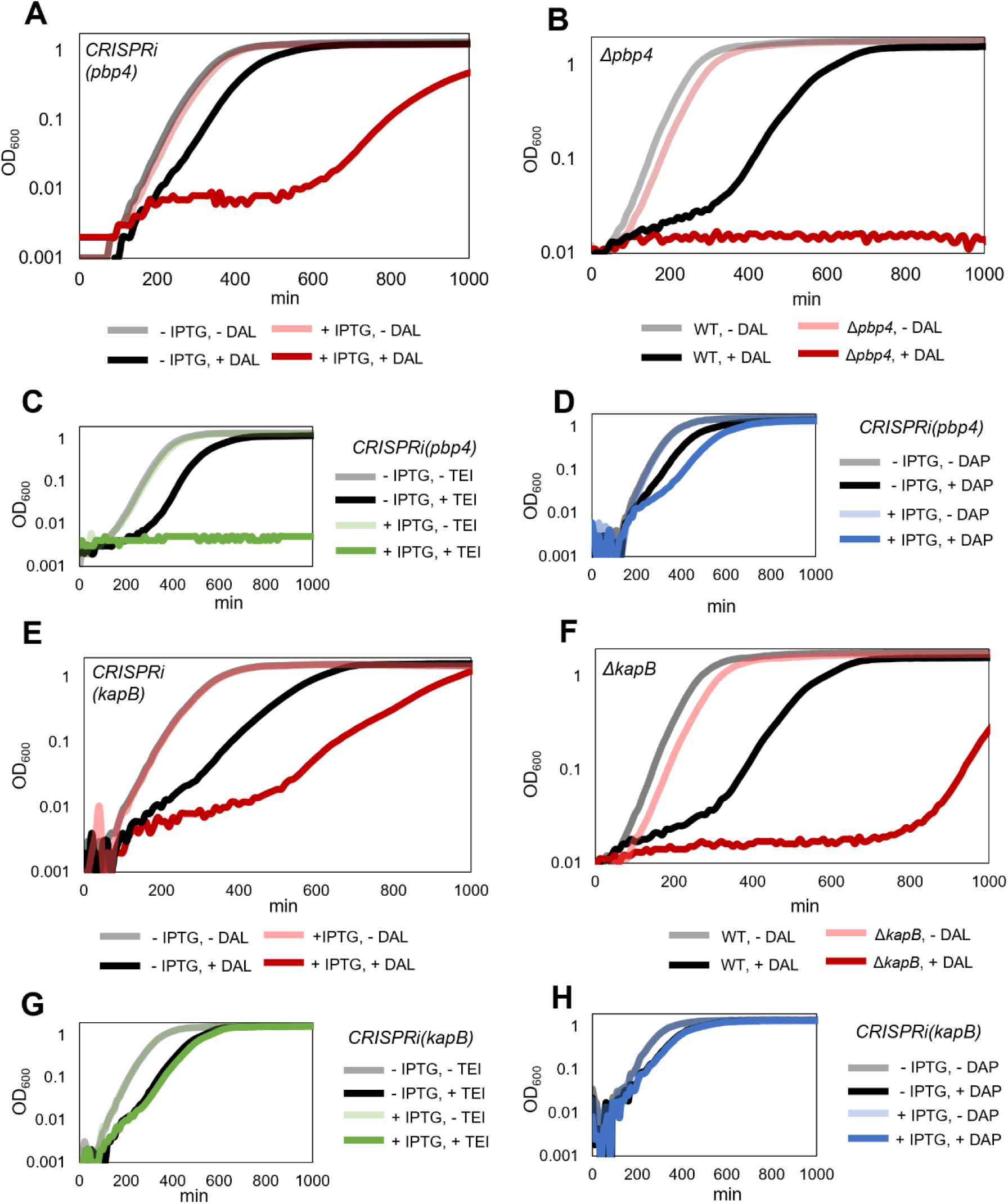
Contribution of *pbp4* and *kapB* to susceptibility of *S. aureus* towards dalbavancin, teicoplanin and daptomycin. (A-D). The presence of functional *pbp4* results in increased tolerance to dalbavancin (A-B), teicoplanin (C) and daptomycin (D). (E-H). *kapB* influences susceptibility to dalbavancin, but not to teicoplanin or daptomycin.

### KapB influences dalbavancin, but not other glycopeptide or lipopeptide antibiotic susceptibility

Several hits identified by the CRISPRi-seq screen have previously been linked to glycopeptide or lipopeptide antibiotic susceptibility like vancomycin and daptomycin. Such hits include the *dltABCD* operon (44), *vraFG* (5), *ezrA* (5, 44), SAOUHSC_00678 (5), and *pbp4* (5). The *dltABCD* operon (**Fig. 4D**) is responsible for modifying cell surface charge by D-alanylation of teichoic acids. The *vraFG* (for vancomycin resistance-associated F and G) operon (**Fig. 4E**) encodes a putative ABC transporter system and was reported as an important factor for vancomycin and daptomycin resistance (5). EzrA is the cell division and septum formation regulator (**Fig. 4F**), and mutants in this gene have previously been shown to be hypersensitive towards daptomycin (5, 44). The uncharacterized gene SAOUHSC_00678 (**Fig. 4G**) has previously been found to be associated with susceptibility to daptomycin, although its function is unknown (5). Finally, a connection between daptomycin or vancomycin sensitivity and *pbp4* has also been reported in some strains (5). To test whether the confirmed hits were general factors affecting susceptibility to glycopeptide or lipopeptide antibiotics, we exposed individual CRISPRi mutants to teicoplanin or daptomycin. Indeed, most of the hits, including *pbp4* (**Fig. 5A, C, D**), also displayed increased sensitivity to antibiotics other than dalbavancin. Exceptionally, knockdown or knockout of *kapB* (“kinase associated protein B”, encoding a non-essential protein of unknown function) led to a significant decrease in tolerance towards dalbavancin, but not towards the other antibiotics (**Fig. 5E, G, H**). These results show that dalbavancin has a unique susceptibility determinant, suggesting its mechanism of action differs from that of related antibiotics.

### Depletion of *ebpS* and SAOUHSC_00659 reduces the susceptibility to dalbavancin

In total 11 hits had a significant positive log_2_ fold-change upon dalbavancin treatment in the CRISPRi-seq experiment (**Table 1, Fig. 4B**), suggesting that depletion of these genes would result in reduced sensitivity. We selected five of the top hits (*ebpS*, *mvaS*, *mvaK2*, *sgtB*, SAOUHSC_00659) for further verification with single depletion strains, and knockdown of two of these targets (*ebpS* and SAOUHSC_00659) was confirmed to increase tolerance to dalbavancin. The strains with depleted SAOUHSC_00659 or *ebpS* grew better than the control in the presence of dalbavancin (**Fig. S5**). We also noted that the depletion of EbpS resulted in higher maximum OD in the stationary phase. The three remaining selected candidate genes did not differ from the control. SAOUHSC_00659 encodes a protein of unknown function, and *epbS* a surface-exposed, integral membrane protein known as elastin binding protein (45). The functions of these two proteins are largely unknown, although EbpS has been reported to bind elastin *in vitro* (46, 47), and has also been suggested to be an important factor for biofilm formation under certain conditions (48). Our study suggests that EbpS and SAOUHSC_00659 are somehow involved in a response to cell envelope stress caused by dalbavancin.

### Direct selection of dalbavancin-tolerant strains reveals that the Shikimate pathway modulates dalbavancin tolerance

Besides the CRISPRi-seq screen, the CRISPRi library holds potential in enrichment and selection studies under specific stresses, apart from next-generation sequencing. As a complementation for the CRISPRi-seq screens of dalbavancin-related factors, we used the CRISPRi library to select for strains with reduced susceptibility to dalbavancin as outlined in **Fig. 6A** (see also methods). The sgRNAs harboured by eight colonies shown to display increased dalbavancin susceptibility were identified by Sanger sequencing (**Fig. 6A**). Five of the strains carried sgRNAs targeting *sagB* (SAOUHSC_01895), two *aroB* (SAOUHSC_01482) and one *vrfA* (SAOUHSC_01192). To verify that the increased resistance was not caused by any secondary mutations in the genome, the sgRNA plasmids were re-introduced into NCTC8325-4 to make new, individual CRISPRi mutants. Dalbavancin susceptibility of these mutants was tested and the results confirmed that knockdown of these genes resulted in reduced dalbavancin sensitivity (**Fig. 6B**).

**Fig. 6.**
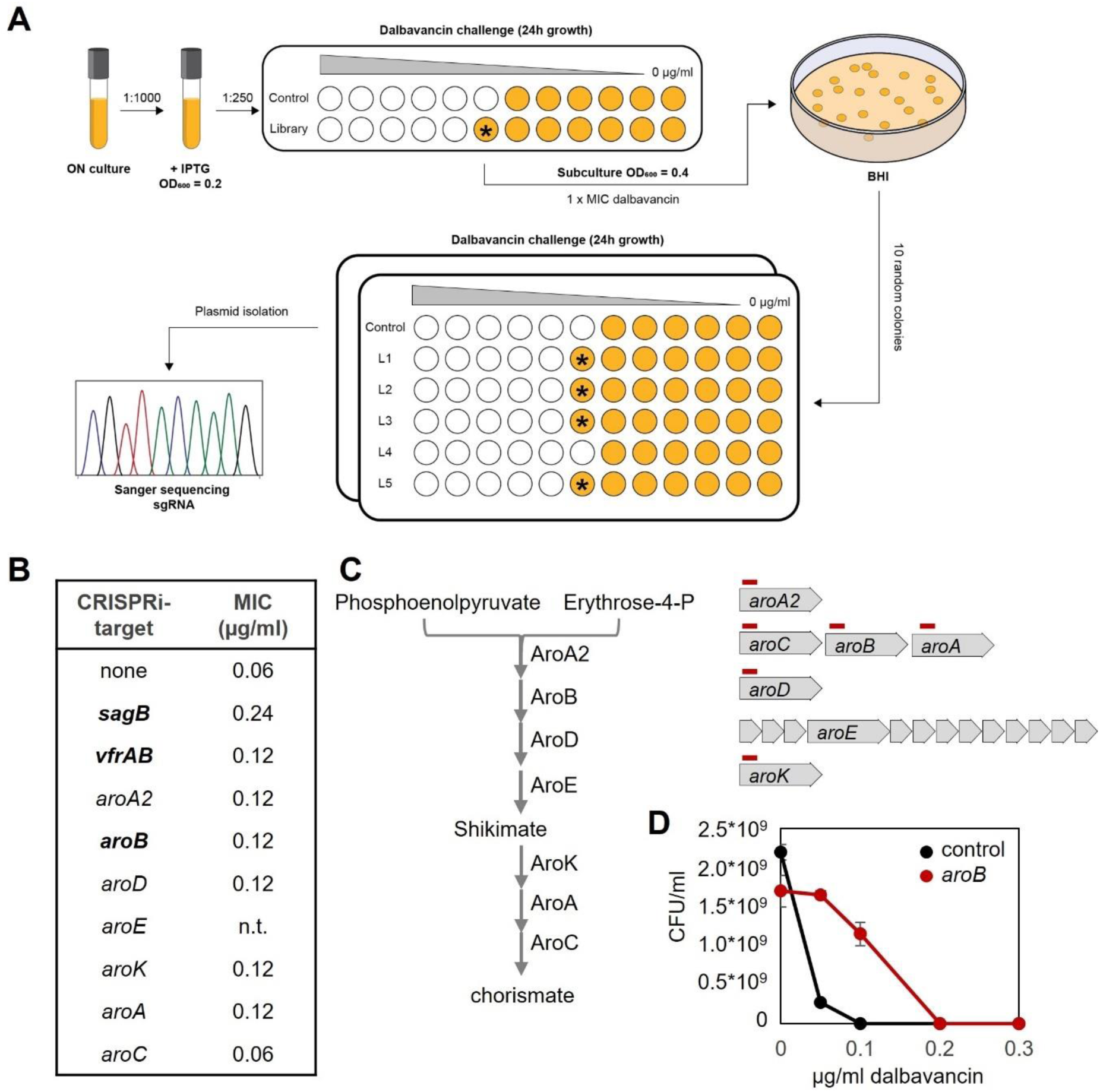
Direct selection of dalbavancin tolerant strains (A) Schematic outline of the CRISPRi library selection study for dalbavancin susceptibility. The asterisks points to wells with increased growth compared to the controls. (B) List of hits for verification studies. The three hits identified by the CRISPRi library selection study are indicated in bold. (C) Schematic illustration of Shikimate pathway (left) and corresponding gene arrangement on the genome (right). The red lines indicate binding sites of the sgRNAs. (D). Population analysis profile showing that the *aroB* knockdown strain survives at higher concentrations of dalbavancin compared to the control.

Knockdown of *sagB* led to a 4-fold increase in MIC to dalbavancin (**Fig. 6B**). *sagB* encodes a glucosaminidase involved in peptidoglycan polymer length control in *S. aureus* (49, 50). Consistent with our discovery, mutations in *sagB* were previously reported to reduce sensitivity to vancomycin due to the extended peptidoglycan polymers present in the mutant (49). The reduced sensitivity to dalbavancin might be caused by the same mechanism. The *vfrA*-gene, whose function is unknown, is encoded on the same operon as *fakA* (formerly known as *vfrB*). FakA encodes a fatty acid kinase which is needed for utilization of exogeneous fatty acids for lipid synthesis. Deletion of *fakA* has been shown to alter the metabolic state of the cell, including changes in intracellular amino acid concentrations (51). AroB encodes the 3-dehydroquinate synthase in the Shikimate pathway for biosynthesis of the amino acid precursor chorismate (**Fig. 6C**), and knockdown of *aroB* was confirmed to increase dalbavancin tolerance (**Fig. 6D**). The link between chorismate biosynthesis and susceptibility to glycopeptides has not been reported before. To further verify these results, CRISPRi strains to knockdown other genes in the Shikimate pathway were constructed: *aroA2*, *aroC*, *aroA*, *aroD*, and *aroK* (**Fig. 6C**). The dalbavancin MIC was determined, and indeed the MIC increased slightly in all strains except for *aroC* (**Fig. 6B**), showing that disruption of the Shikimate pathway modulates susceptibility to dalbavancin in *S. aureus* NCTC8325-4.

## Discussion

In this work we have developed a pooled, genome-wide, compact operon-based CRISPRi library with 1928 sgRNAs for *S. aureus* (**Fig. 2**). To enable high-throughput genetic screening, we combined the CRISPRi library with Illumina sequencing and developed CRISPRi-seq in *S. aureus*. In contrast to the previously developed *S. aureus* CRISPRi library, which was made by a CRISPR adaptation strategy and contains an average of 100 guide RNAs per gene (22), the compactness of our library with one systematically designed sgRNAs per gene requires less sequencing depth and could more easily overcome bottlenecks in CRISPRi-seq experiments, which may occur in some experimental setting such as in animal models (16). Initially, CRISPRi-seq was employed to quantify the essentialome in two *S. aureus* strains, NCTC8325-4 and Newman (**Fig. 3**). By comparative analysis of the essentialomes of these two strains by CRISPRi-seq with published essentialome data, we showed that CRISPRi-seq could capture the conserved essential genes across different *S. aureus* strains. Approximately 75% of the sgRNAs expected to target conserved essential genes according to transposon-based knockout studies in related *S. aureus* strains (40–42) were indeed defined to target essential genes in the CRISPRi-seq screen. Nevertheless, a significant fraction (19%) of the conserved essential genes was not defined as essential in our CRISPRi-seq results. This may be explained by different strains used and differences in growth conditions between the studies: gene essentiality is strain-and condition-specific. Alternatively, some of these missing hits may represent false negatives, resulting from sgRNAs with poor efficiency or insufficient depletion during the experiments. Lastly, differences between studies may be partially explained by differences in the definition of essentiality, as in our studies we used relatively strict statistical parameters for this designation (significantly more than doubling or halving of the CRISPRi strain abundance upon CRISPRi induction), and transposon mutagenesis-based studies requires different statistical approaches to determine essentiality.

It is expected that the number of essential genes would increase with the number of generations of growth during CRISPRi induction. Two independent CRISPRi-seq experiments (12- and 20-generation of growth) were performed in this study, however, there were very few differences between the two experiments (**Fig. 3**). Thus, 12 generations of growth are sufficient to capture the CRISPRi-seq essentialome in these experimental conditions. Furthermore, our *in silico* analysis suggests that the sgRNA library is largely functional in other, commonly used *S. aureus* strains (64-89% of sgRNAs with an exact non-template strand within-gene binding site), covering most of their annotated genetic elements either directly, by binding to the annotated gene (53-63%), or indirectly, by polar effects (79-93%) (**Fig. S1, Table S4-S10**). Note that the former percentages are based on actual annotations, i.e., they are measured, whereas the latter are estimated by extrapolation, assuming similar operon structures between all strains and NCTC-8325-4 (as defined by Mäder et al. (2016)). It is possible, however, that these estimates are still on the conservative side, as differences in annotations might hinder target gene detection and sgRNAs might still be active at sites with minor allelic differences between homologs, as long as the mismatches are few and on the PAM-distal side of the sgRNA spacer. Although we have indeed shown the library’s functionality in one such strain, Newman (**Fig. S4**), it should be noted that its use in any other strain will be inferior to NCTC8325-4, which the library was designed for.

We next explored genes related to dalbavancin susceptibility using CRISPRi-seq and CRISPRi-oriented selection studies (**Fig. 4** and **Fig. 6**). Dalbavancin is a semi-synthetic lipoglycopeptide with broad activity against Gram-positive bacteria, including *S. aureus* MRSA strains (25). To get further insights into the mechanism of action of dalbavancin and its unique features, we applied CRISPRi-seq under dalbavancin stress. Interestingly, we identified and validated both genes that led to increased and decreased dalbavancin tolerance. We found many genes known to be involved in susceptibility to vancomycin (glycopeptide), teicoplanin (vancomycin-derivative) or daptomycin (lipopeptide) to also influence dalbavancin sensitivity. This indicates that there are similar mechanisms contributing to resistance and sensitivity for these antibiotics, which is unsurprising given that they all target the cell wall and/or membrane.

However, we also showed that a new factor, *kapB*, encoding a 127 aa conserved protein, specifically contributes to *S. aureus* susceptibility towards dalbavancin, but not other glycopeptides or lipopeptides (**Fig. 5**), suggesting that dalbavancin has an antibacterial mechanism which to some extent differs from teicoplanin and related antibiotics. The function of KapB and how it affects dalbavancin sensitivity remains to be fully explored. *kapB* is monocistronic and located between the *mnhABCDEFG* operon and *ppi*, encoding a multicomponent monovalent cation/H+ antiporter system and a putative peptidyl-prolyl cis-trans isomerase, respectively. Knockdown of neither of these neighbouring genes had any significant impact on dalbavancin susceptibility in the CRISPRi-seq screen, suggesting that KapB acts independently (**Table S17**).

Furthermore, in our assays, *S. aureus* NCTC8325-4 mutants lacking PBP4 became more susceptible to dalbavancin and teicoplanin (**Fig. 5**). PBP4 is a transpeptidase in *S. aureus* important for the formation of cross-links in *S. aureus* peptidoglycan. In contrast to our findings here, inactivation or reduced expression of *pbp4* have previously been linked to glycopeptide-resistant mutants, which has suggested a mechanism where resistance is caused by the accumulation of non-crosslinked muropeptides in these mutant strains, resulting in binding and sequestering of the antibiotics (52, 53). However, in contrast to the NCTC8325-4 strain, glycopeptide-resistant mutants often carry multiple, diverse mutations contributing to the resistance phenotype (35, 54). Moreover, previous experiments with Δ*pbp4* mutants in glycopeptide-sensitive strains showed that the vancomycin MIC is the same as for their respective wild-type (53, 55, 56). We also confirmed that this was the case for NCTC8325-4 (MIC of 2 µg/ml vancomycin for both wild-type and Δ*pbp4*). In addition, it was recently reported that a *pbp4* deletion increased the susceptibility to vancomycin in a bone tissue infection model (56). The association between PBP4 and glycopeptide antibiotics in different genetic backgrounds should therefore be further investigated, particularly since dalbavancin has been shown to act synergistically with different β-lactam antibiotics (57–60).

Knockdown of either of two essential genes, *ebpS* and SAOUHSC_00659, increased tolerance to dalbavancin in the CRISPRi-seq screen. Note that these genes could not be identified by any knockout-based screening due to their essentiality for growth, demonstrating that CRISPRi-seq can be used as an important complementary technique for high-throughput screenings in *S. aureus*. The molecular mechanisms explaining why EbpS and SAOUHSC_00659 influence dalbavancin susceptibility remain to be elucidated.

We also used a direct selection approach to screen for strains with reduced sensitivity to dalbavancin (**Fig. 7**), and found other genes (*sagB*, *aroB* and *vfrAB*) than by CRISPRi-seq (**Fig. 4**), demonstrating that the output of such screens depends on the experimental setup. Indeed, the hits identified in the direct selection approach were not significantly differentially essential following the CRISPRi-seq approach (**Table S17**). Of these hits, SagB, a glucosaminidase cleaving strands of peptidoglycan, has previously been shown to result in reduced vancomycin sensitivity (49). The *vfrA-fakA* operon encodes proteins with similarity to acid shock protein Asp23 and a fatty acid kinase, respectively. The role of VfrA is unknown, but previous studies have shown that FakA is responsible for incorporation of exogeneous fatty acids into phospholipids *in vivo* (61). Notably, mutations in *fakA* have indeed been identified in dalbavancin-resistant mutants generated by antibiotic exposure (36). Of note, we also demonstrate that knocking down the Shikimate pathway can result in reduced sensitivity to dalbavancin *in vitro* (**Fig. 6**). The Shikimate pathway has been considered a promising target for antibacterial drugs, because it has no counterpart in mammals and is essential for bacterial growth and virulence in many contexts (62). However, the results shown here argue against Shikimate pathway genes as therapeutic targets, as their depletion may also reduce antibiotic susceptibility.

In this work, we have created a tight, compact CRISPRi library and showed its functionality and utility in *S. aureus* in both CRISPRi-seq and direct selection experiments. We have uncovered genes previously unknown to be involved in antibiotic susceptibility, among which core essential genes which cannot be detected using more traditional mutagenesis approaches as such mutants are by definition excluded from those libraries due to their inability to grow. This study will serve as a steppingstone to explore the molecular mechanisms involved in antibiotic tolerance in general, and of dalbavancin in particular.

## Materials and methods

### Bacterial strains and growth conditions

*Staphylococcus aureus* strains were grown in brain heart infusion (BHI) or tryptic soy broth (TSB) medium at 37 °C with shaking or on BHI agar plates. When needed, 5 µg/ml erythromycin and/or 10 µg/ml chloramphenicol were added for selection. Transformation of *S. aureus* was performed with electroporation as described before (63). *Escherichia coli* was grown on LA agar or in LB medium at 37 °C with shaking. Ampicillin (100 µg/ml) was added for selection. *E. coli* was transformed using heat shock of chemically competent cells. A list of all strains used in the study is found in **Table S18**.

### Plasmid and strain construction

#### pLOW-P_spac2_-dcas9

The P*_spac_* promoter of the pLOW-plasmid used to construct our original CRISPRi system contains a single *lacO* operator site (12, 37). An extra *lacO* operator site was introduced 58 nt upstream of the pre-existing *lacO*-operator, to create the promoter P*_spac2_*. The position of the *lacO* site was selected based on the P_spac_ promoter in plasmid pLA8-1, which is tightly regulated with a high dynamic range in *Streptocccus pneumoniae* (64). The *lacO* site was introduced by inverse PCR, using primers mvh54_Pspac_Rev and mvh55_intro_lacO_Pspac with pLOW-*dcas9* as template. The product was digested with DpnI and transformed into *E. coli* IM08B. Correct transformants were verified by PCR and sequencing. The primers used in this study are listed in **Table S19**.

#### pVL2336

The plasmid expressing the sgRNAs, pCG248-sgRNA(*xxx*) (12), was modified to allow Golden Gate cloning and library preparation for Illumina sequencing. The two BsmBI sites on the vector were removed by circular PCR with OVL2127 and OVL2128 followed by BsmBI digestion and T4 self-ligation. After that, the vector pCG248-sgRNA(*xxx*) was linearized by digestion with BglII and BamHI; and a DNA fragment containing read1 (Illumina sequencing element), P3 (constitutive promoter for sgRNA expression), BsmBI site (producing cut-edge at +1 of P3), mCherry (as reporter for the cloning), second BsmBI site (producing cut-edge between the base pairing region and dCas9 handle binding region) and read2 (Illumina sequencing element) was amplified from pPEPZ-sgRNAclone (Addgene# 141090) with OVL2152 and OVL2153 (23). The amplified DNA fragment was then cloned into pCG248-sgRNA(*xxx*) to replace the pCG248-sgRNA(*xxx*) fragment by infusion cloning, resulting in the sgRNA cloning vector pVL2336.

#### Construction of single sgRNA plasmids

New 20-bp sgRNA sequences were cloned into pVL2336 using Golden Gate cloning as described before (16). Briefly, two oligos for each sgRNA (**Table S20**) were diluted in TEN buffer (10 mM Tris, 1 mM EDTA, 100 mM NaCl, pH8), incubated at 95°C for 5 min and then slowly cooled down to room temperature for annealing. The vector pVL2336 was digested with BsmBI to remove the mCherry fragment and purified by gel extraction. The purified vector and the annealed oligos were ligated using T4 ligase and transformed into *E. coli* IM08B. Correct vectors were verified by Sanger sequencing with primer MK25.

### Construction of deletion mutants (Δ*pbp4*::*spc* and Δ*kapB*::*spc* mutants) in *S. aureus* NCTC8325-4

pMAD-pbp4::spc was assembled using overlap extension PCR and ligated into pMAD using restriction cloning. The region upstream and downstream of *pbp4* were amplified with primer pairs mk411/mk412 and mk413/mk414, respectively. The spectinomycin resistance cassette was amplified with primer pair mk188/mk189. The inner primers mk412 and mk413 contain overlapping sequences, allowing fusion of the fragments in a second step PCR to make the fusion construct *pbp4*_up-*spc-pbp4*-down. The fused fragment was digested with NcoI and BamHI, whose restriction sites were introduced with the outer primers mk411 and mk414, respectively, and ligated into the corresponding sites of pMAD. The ligated plasmid was transformed into *E. coli* IM08B and verified by PCR and sequencing.

pMAD-kapB::spc was assembled and ligated into pMAD-GG (pMAD-plasmid adapted for Golden Gate cloning, laboratory collection) by Golden Gate cloning. The up- and downstream regions were amplified with primer pairs AHF13/AHF14 an AHF17/AHF18, respectively. The spectinomycin cassette was amplified with AHF15/AHF16. BsaI restriction sites generating complementary overhangs are introduced in the primers. The purified fragments and the pMAD-GG vector were digested with BsaI and assembled using NEB Golden Gate Assembly kit. The assembly reaction was transformed into *E. coli* IM08B and verified by PCR and sequencing.

Following construction of the plasmids, the deletion strains were made as previously described for the temperature sensitive pMAD system (65). Plasmids were transformed into *S. aureus* at permissive temperature (30°C) and plated onto X-gal plates with erythromycin selection. Blue colonies were re-streaked at 30°C and verified by PCR. A single colony was picked in medium without selection and incubated at 30°C for 2 h before the tube was transferred to 43°C for 6 h. The culture was plated onto TSA plates with X-gal and spectinomycin for selection and incubated at 43°C. Candidates for double crossover (white colonies), were re-streaked on two separate plates to identify colonies that were spectinomycin resistant and erythromycin sensitive, and strains were further verified by PCR and sequencing.

### CRISPRi library construction

#### Design of sgRNAs

The coding sequences of the selected genes were used as input for PAM motif search using CRISPR Primer Designer (Yan et al., 2015). The 20-bp sequence adjacent to the 5’ proximal PAM, targeting the non-template strand was selected as candidate sgRNA target sequence. To check the specificity of the 20-bp sequences, the core 12-bp sequence adjacent to the PAM (core sgRNA) was used for a BLAST search, using the NCTC8325 genome as target, to filter out non-specific targets. The 20-bp sequence closest to the 5’ end of the gene fulfilling these criteria was selected as the final target. Finally, 1928 sgRNAs were designed (**Table S1 and S20**)..

#### *Construction of the genome-wide sgRNA library in E. coli* IM08B

The 20-bp base-pairing region of the sgRNAs was first cloned into pVL2336 in *E. coli* IM08B by Golden Gate cloning. 1.3×10^6^ transformant colonies were obtained, providing a 674-fold coverage of the 1928 sgRNAs library. The false positive ratio of sgRNA cloning was estimated by calculating the percentage of the red colonies among all transformant colonies, which showed a false positive rate lower than 0.03% (no red colony identified among 3000 colonies). The plasmids carrying the sgRNAs were then purified from *E. coli* IM08B, and used as template in the one-step PCR reaction for Illumina amplicon library preparation as previously described (23).

### Construction of CRISPRi library in *S. aureus* NCTC8325-4 and Newman

Plasmids isolated from the *E. coli* sgRNA library were transformed into *S. aureus* MH225 (NCTC8325-4, pLOW-Pspac2-dcas9) and *S. aureus* MH226 (Newman, pLOW-Pspac2-dcas9) by electroporation. Multiple parallel transformation reactions were plated onto BHI agar plates containing 5 µg/ml erythromycin and 10 µg/ml chloramphenicol. More than 200 000 colonies were collected using a cell scraper, pooled and resuspended in BHI containing 5 µg/ml erythromycin and 10 µg/ml chloramphenicol. The pooled libraries were diluted to OD600 = 0.2 in the same medium, re-grown to OD_600_ = 0.8 and stored as glycerol stocks for further use.

### CRISPRi-seq essentiality screen

Growth of the library for CRISPRi-seq experiments was performed in 100 ml BHI medium supplemented with 5 µg/ml erythromycin and 10 µg/ml chloramphenicol for selection using 250 ml Erlenmeyer flasks. Cultures were grown with shaking at 37°C. When appropriate, 250 µM IPTG was added for induction. Four parallels per condition were used. For the 20-generation experiment, the library was inoculated 1/1000 from the stock, grown until OD_600_ = 0.8, then reinoculated 1/1000 in fresh medium and again grown until OD_600_ = 0.8. The cultures were then transferred to 50 ml Nunc tubes and collected by centrifugation (6000 x g, 5 min, 4°C). For the 12-generation experiment, the library was inoculated 1/4000 from the stock and grown to OD_600_ = 0.8 before cells were harvested. For the dalbavancin experiment, the same conditions as in the 12-generation experiment were used, except that a sublethal concentration (0.015 µg/ml, equivalent to ¼ of the MIC) was added. For plasmid isolation, cells were lysed by treatment with lysostaphin (40 µg/ml, 30 min at 37°C) using QIAGEN Plasmid Midi Kit.

### Library preparation and Illumina sequencing

For the 12 generation experiment, construction of the amplicon library for Illumina sequencing was performed as described in our previous study (de Bakker, 2022). Specifically, plasmids were purified from the *S. aureus* CRISPRi library samples. Concentration of the plasmids was quantified with Nanodrop. To prepare the Illumina amplicon library, 1 µg of plasmids was used as template for the one-step PCR with primers described before (de Bakker, 2022). The PCR products were purified by gel extraction. The purified products were then quantified by Qubit and processed for MiniSeq sequencing with a custom recipe as described previously (23).

For the 20-generation experiments, the Illumina amplicon library to be sequenced was amplified using the condition described previously (66). Briefly, 2 μl of the plasmid was used as a template and amplified using the Nextera DNA Indexes (Illumina Inc., San Diego, CA, USA) index primers. The PCR product was then purified and normalized using SequalPrep Normalization Plate Kit (Thermo Fisher Scientific, Waltham, MA, USA). The purified library was quantified using the KAPA library quantification kit (KAPA Biosystems, Wilmington, MA, USA) and sequenced on an Illumina Miseq platform (Illumina Inc) using the Miseq Reagent Kit v3 (Illumina Inc.).

### CRISPRi-seq differential enrichment analyses

Read pairs of the paired-end sequencing data of the NCTC8325 20-generation experiment and the sequenced plasmid pool were merged using PEAR (v0.9.11) (67) prior to sgRNA count extraction with 2FAST2Q (v2.5.0) (68) as for the other (single-end) sequencing data. Because of poor sequencing quality on the read extremes for the single-end sequencing data, alignment with 2FAST2Q was performed using the 2-17 nucleotides of the sgRNA sequences for those files. Otherwise 2FAST2Q was used with default parameters. sgRNA depletion/enrichment was tested using DESeq2 in R (v4.1.1) (69). We always tested against an absolute log_2_FC of 1 at a significance level of α=0.05. For Principal Component Analyses, counts were normalized with DESeq2’s blind rlog transformation. Sample 9_S9_L001_R1_001 (screening for NCTC8325, 12 generations, uninduced, 1^st^ replicate) showed aberrant read count patterns and did not correlate with its replicates, so was excluded from further downstream analyses. For comparison of 12- and 20-generation experiments, log_2_FC values were scaled, but not centered, through division by the root mean square of the values per experiment using the built-in scale() function in R.

### Evaluation of target efficiency of the sgRNA library

sgRNA library efficiency on several genomes was evaluated using a previously published pipeline: https://github.com/veeninglab/CRISPRi-seq (23). The script *sgRNA_library_evaluation_cmd.R* was run with default settings, and genbank files as input. Genbank files with the following nine assembly accession numbers were obtained from the NCBI RefSeq database, accessed on 5-SEP-2022: GCF_000009005 (RF122), GCF_000009665 (Mu50), GCF_000010465 (Newman), GCF_000011505 (MRSA252), GCF_000012045 (COL), GCF_000013425 (NCTC8325), GCF_001027105 (DSM20231), GCF_002085525 (JE2-USA300) and GCF_900475245 (NCTC8325). The total numbers of annotated features (i.e., unique locus_tag flags) were extracted from the genbank files (2743, 2910, 2930, 2881, 2792, 2872, 2761, 2881 and 2858, respectively) and used in combination with the pipeline summary output files to compute the percentage of features that were covered by direct binding of at least one sgRNA within the feature boundaries on the non-template strand, without mismatches. Indirect coverage was then estimated by extrapolation: according to the summary output file, 1864 NCTC8325 (GCF_000013425) features were directly targeted. We know from the design phase that 2764 of 2836 features were targeted considering known operon structures and polar effects. We then estimated for each genome the number of indirectly targeted features as the number of direct targets multiplied by 2764/1864, assuming similar operon structures among the strains. Note that due to differences between the current RefSeq annotations and those used in the design phase, according to this estimate the target genome is covered 96.2%, instead of the pre-computed 2764/2836=97.5%. Similarly, library functionality was computed as the percentage of sgRNAs that have a zero-mismatch binding site on the non-template strand within the boundaries of an annotated feature in each genome.

### Growth assay

Growth curves were monitored in 96-well microtiter plates. Overnight cultures were diluted 100-fold in fresh medium, and then 200 µl of the diluted culture was added into each well of 96-well plate as the starting culture. The microtiter plates were incubated at 37°C and OD_600_ was measured every 10 minutes using Synergy H1 Hybrid Reader (BioTek) or Hidex Sense (Hidex Oy). When appropriate, erythromycin and chloramphenicol were added for selection and IPTG was added for induction.

### Antibiotic susceptibility assays

Susceptibility to antibiotics (dalbavancin, teicoplanin, and daptomycin) in liquid medium was determined using two-fold dilution assays in microtiter plates. Two-fold dilution series of the antibiotics were made in growth media containing erythromycin and chloramphenicol (for selection). When appropriate, IPTG was added for induction. Unless otherwise mentioned, 250 µM IPTG was used. For the dalbavancin assays, 0.002 % Tween-80 was added to the medium. For daptomycin, the assays were performed in the presence of 0.05 mg/ml CaCl_2_. Exponentially growing cells were diluted 100-fold. The microtiter plates were incubated at 37°C and OD_600_ was measured every 10 minutes using Synergy H1 Hybrid Reader (BioTek) or Hidex Sense (Hidex Oy). All assays were repeated at least twice.

### Population analysis profile (PAP)

PAPs were performed as described (70). Briefly, cultures induced with 250 µM IPTG were grown to OD 0.8 and diluted 10^−4^ to 10^−7^. Dilutions were then plated on BHI plates containing dalbavancin (0, 0.1, 0.2, 0.3 and µg/ml). Plates were incubated at 37°C, and colonies were counted after 24 hr.

### Selection of dalbavancin-tolerant mutants

Overnight cultures of the CRISPRi library and a control strain (AHF1010; pLOW-dcas9, pVL2336-sgRNA(nontarget)), were rediluted 1:1000 in BHI containing antibiotics for selection and 250 µM IPTG for induction and grown until OD_600_ = 0.2. Then, a two-fold dilution of dalbavancin (starting from 4 µg/ml dalbavancin) was set up, 250-fold dilutions of the cultures were added and the plates were incubated overnight. From the library, cells from the well with the highest concentration of dalbavancin allowing growth of the library were further inoculated in medium containing 250 µM IPTG and 1хMIC dalbavancin (0.06 µg/ml) until OD = 0.4. From this culture, dilutions were plated onto TSB plates containing chloramphenicol and erythromycin to obtain single colonies. From two independent experiments, ten random colonies were picked and another microtiter assay was performed to determine whether the susceptibility to dalbavancin differed from the control. For strains with increased susceptibility, plasmids were isolated and the sgRNA was sequenced by Sanger sequencing using primer MK25 to identify which gene was depleted.

## Acknowledgments

This work was supported by a Joint Programming for Antimicrobial Resistance grant to MK and JWV from Research Council of Norway (grant 296906) and Swiss National Science Foundation (grant 40AR40_185533). XL was supported by The Science and Technology Project of Shenzhen (JCYJ20220818095602006), National Nature Science Foundation of China (82270012), and Shenzhen University 2035 Program for Excellent Research (86901-00000216).

## Data availability

All sequencing data generated in this study are available on SRA, accession number PRJNA994855.

## Supplemental figures and tables

**Fig S1.**
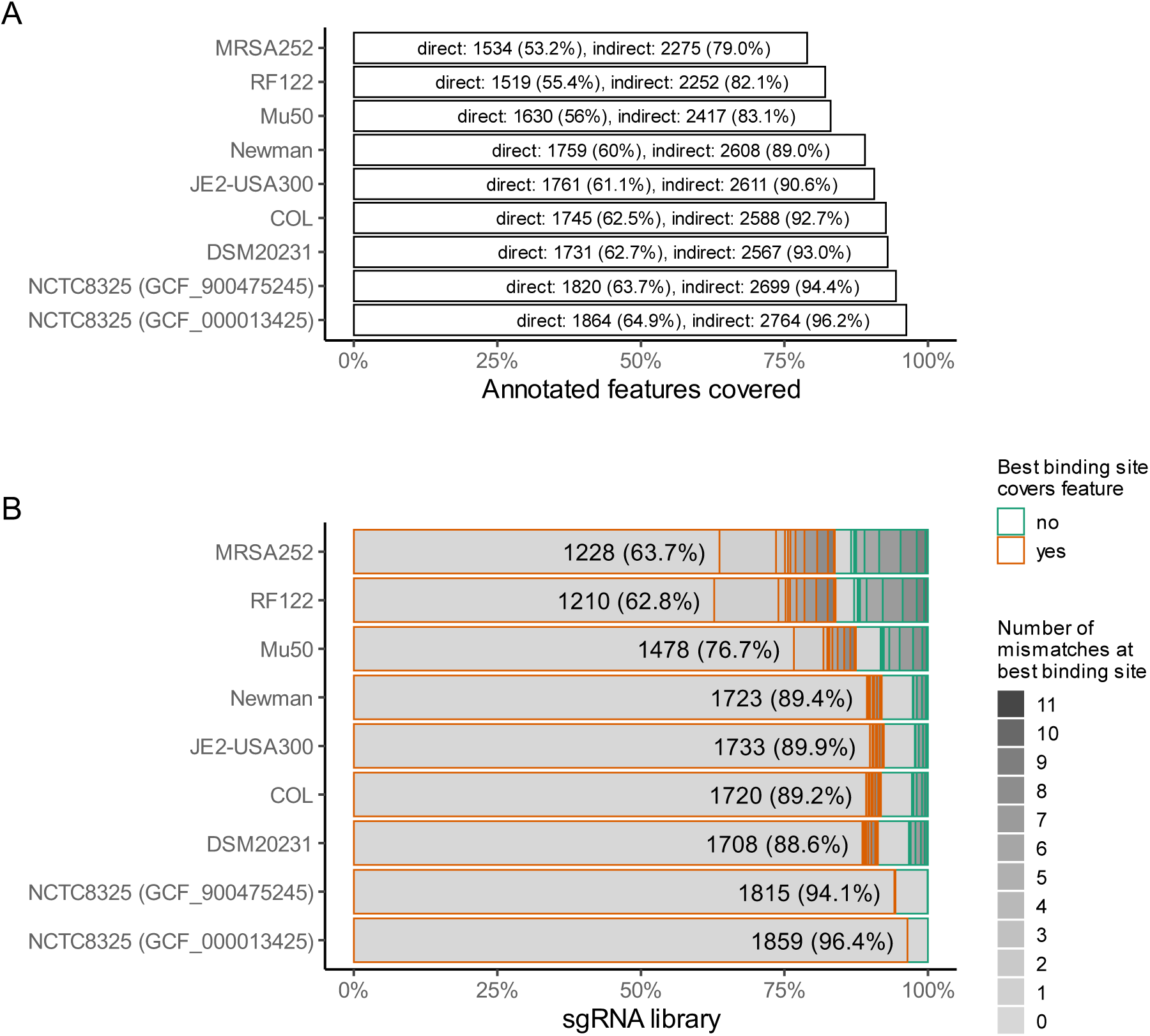
Library coverage across *S. aureus* genomes. (A) Genome coverage of the NCTC8325-based sgRNA library. Overview showing to which extent the library covers the genomes of various *S. aureus* strains. “Direct” indicates the number of features targeted by at least one sgRNA within the feature on the non-template strand, without mismatches. “Indirect” indicates the estimated number of features targeted, accounting for polar effects and operon structures. (B) Overview showing the proportion of the 1928 sgRNAs that are functional in various *S. aureus* strains. Note that two different annotations of the NCTC8325 genome are included (GCF_000013425 and GCF_900475245). For each sgRNA, the best binding site in the genome was identified using the library evaluation pipeline (de Bakker et al, 2022). Bars indicate whether these binding sites are located within annotated features on the non-template strand, and the number of mismatches between the binding site and the sgRNA. Numbers are shown explicitly for sgRNAs targeting an annotated feature without mismatches. Any discrepancies between the actual target genome coverage/library functionality and the estimates reported here are due to discontinued and merged annotations in the current RefSeq version (GCF_000013425).

**Fig. S2.**
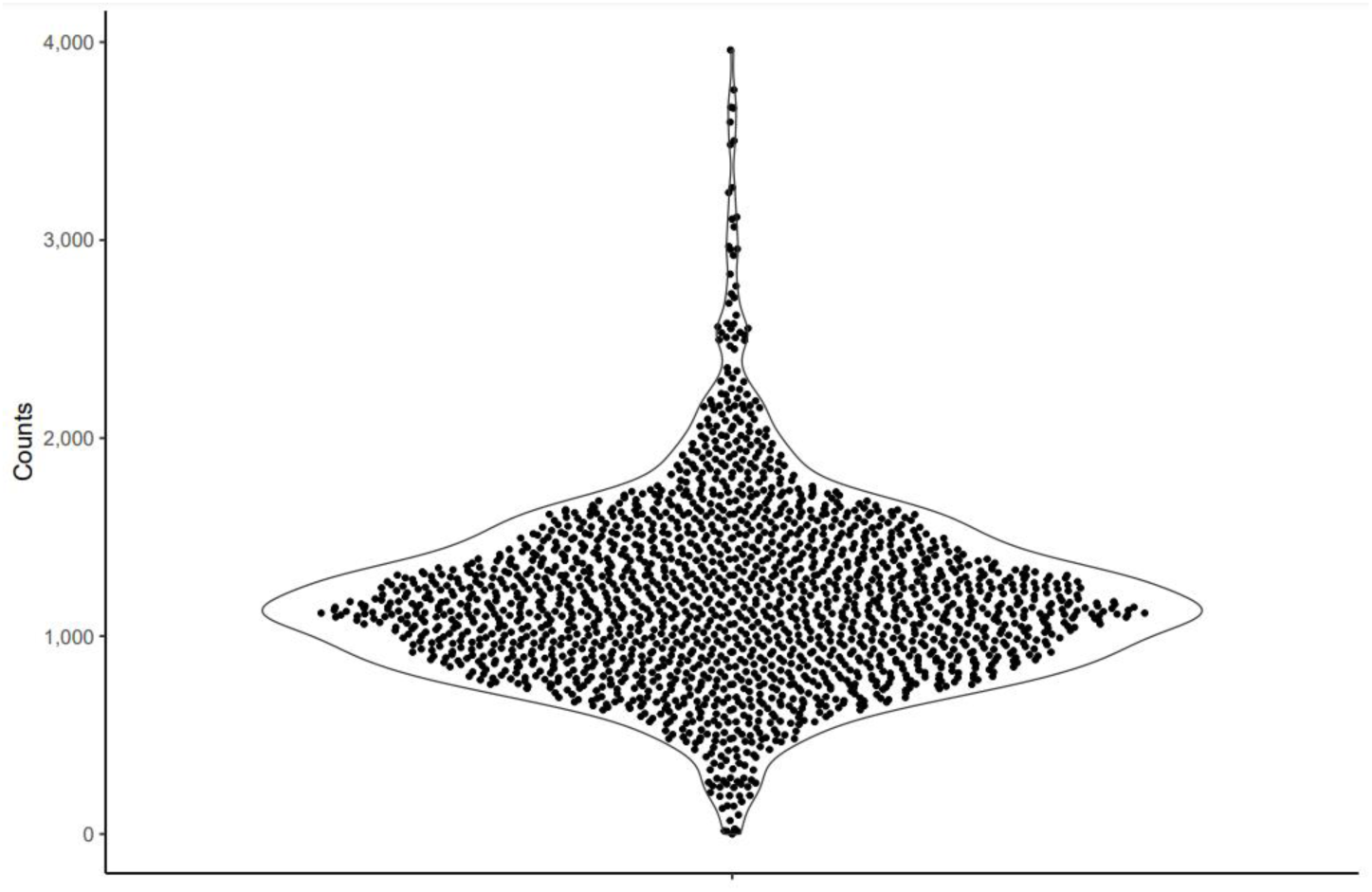
Violin plot showing sgRNA distribution of the plasmids isolated from *E. coli* IM08B. Each dot represents one sgRNA.

**Fig. S3.**
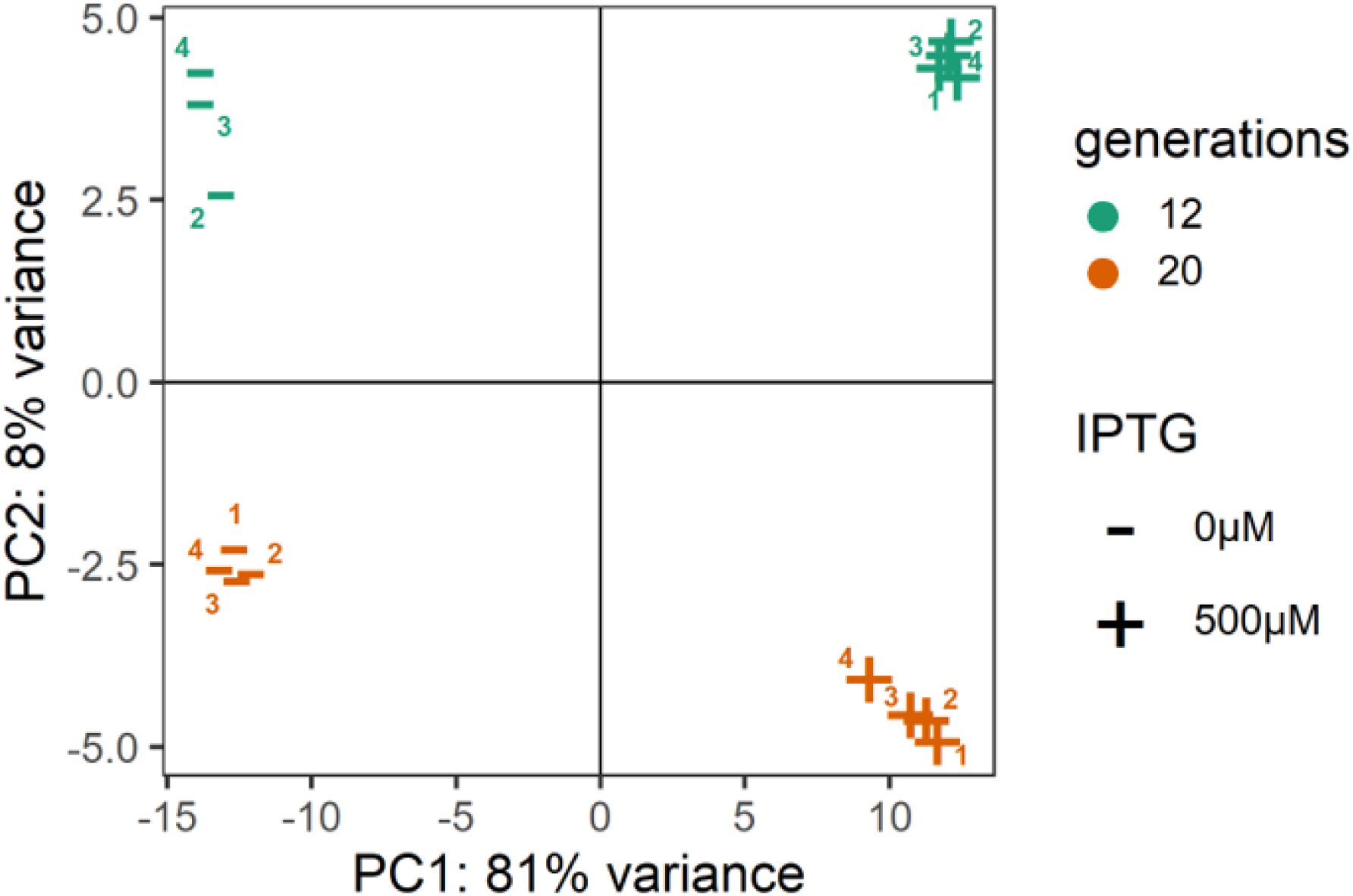
Principal Component Analysis of the rlog-transformed (69) sgRNA counts. Replicate 1 of the uninduced 12-generation experiment samples was excluded, as many CRISPRi strains were depleted from it (without induction), yielding low correlations with the other replicates in the process.

**Fig. S4.**
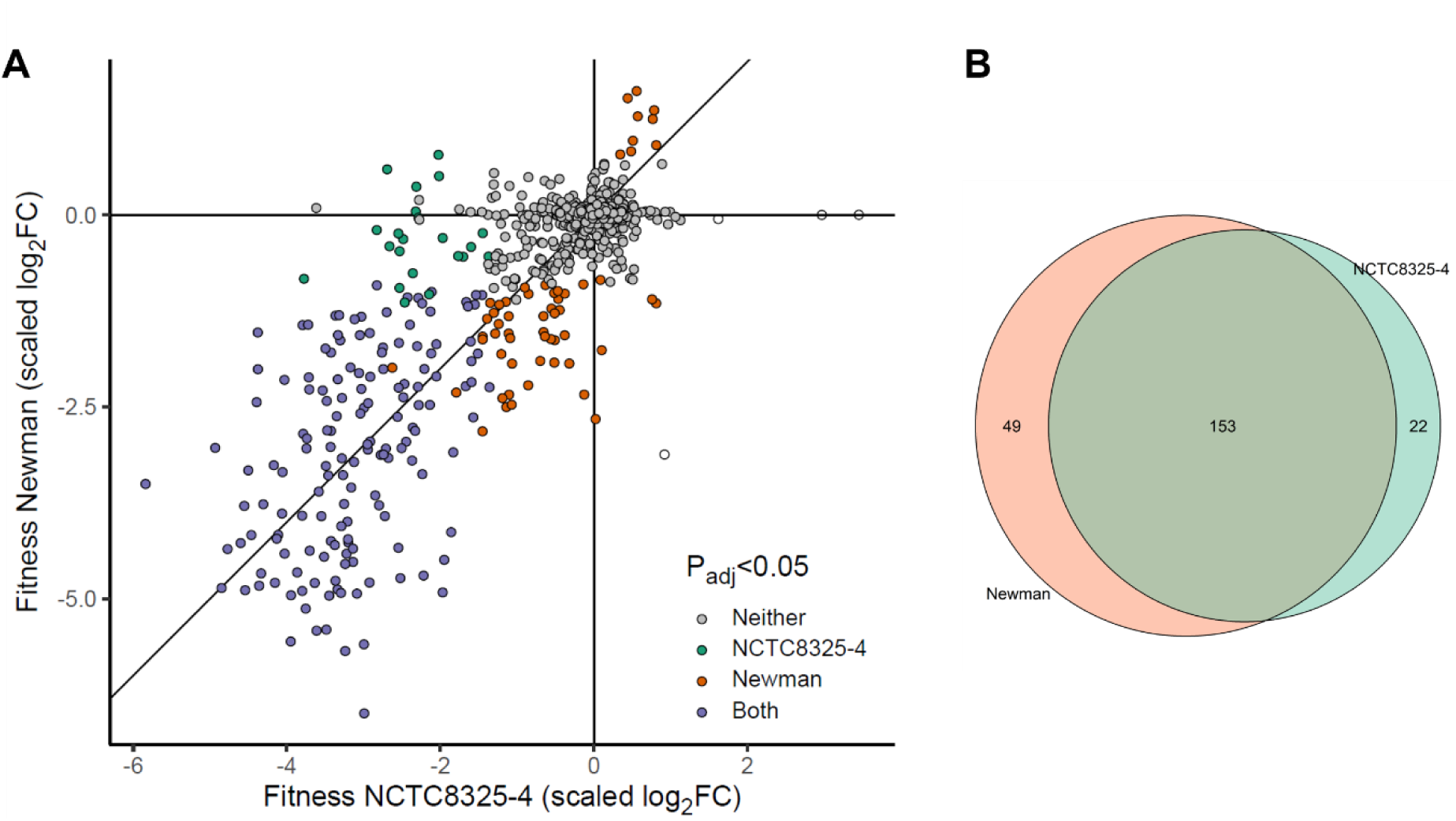
CRISPRi-seq analysis of *S. aureus* Newman grown for 12 generations. (A) Genome-wide fitness comparisons between Newman and NCTC8325-4. The colors indicate sgRNAs that are significantly depleted or enriched in both strains (purple), neither (grey), only in Newman (orange) or only in NCTC8325-4 (green). Four sgRNAs lack color since their NCTC8325-4 p-value could not be computed due to count outliers as detected by DESeq2. (B) Venn diagram showing the overlap in essentialome determined by CRISPRi-seq in *S. aureus* Newman and NCTC8325-4.

**Fig. S5.**
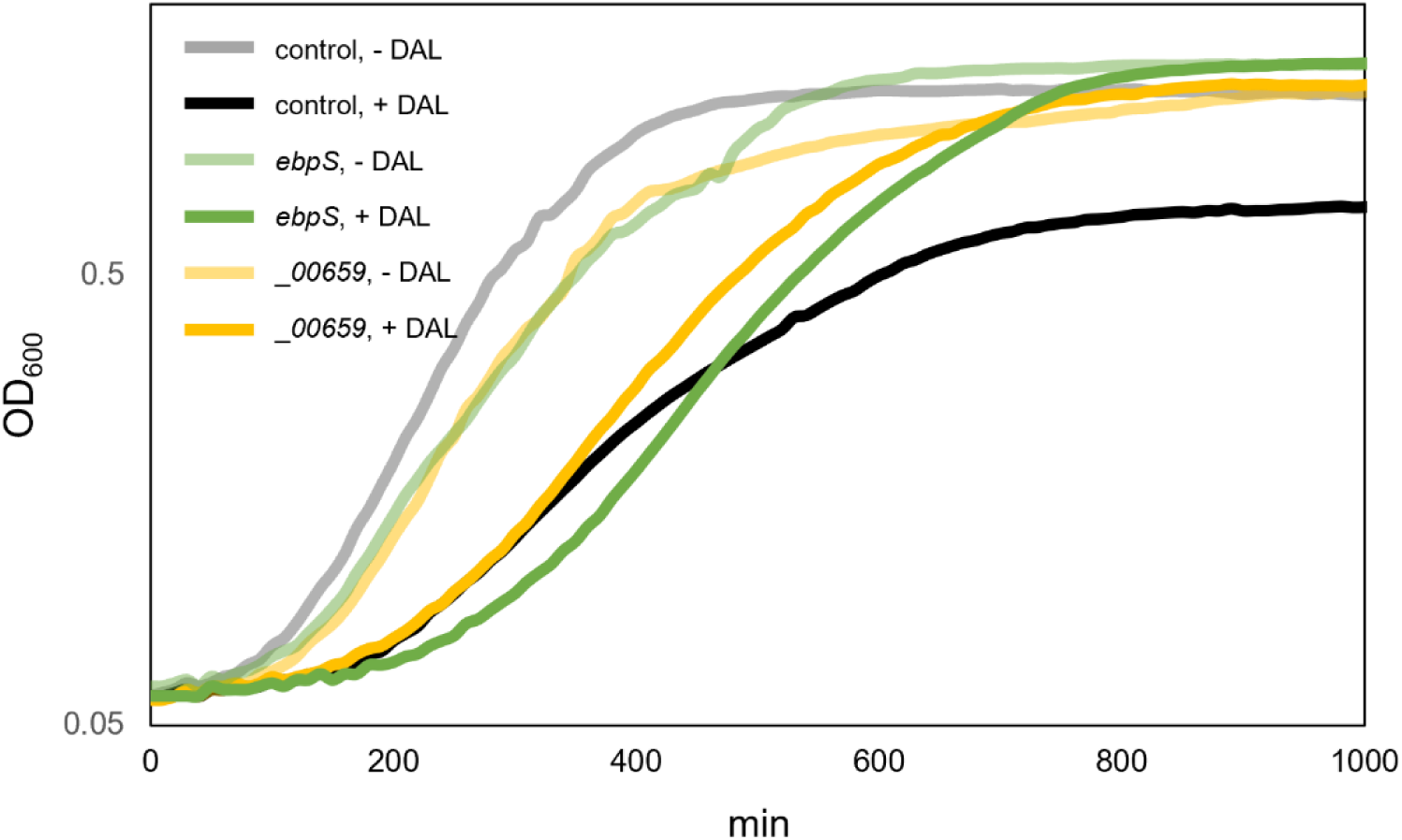
Knockdown of *ebpS* and SAOUHSC_00659 led to reduced susceptibility to dalbavancin. CRISPRi control strain (no-target control, MM75), and CRISPRi strains targeting e*bpS* and SAOUHSC_00659 were all grown in the presence of 250 µM IPTG for induction of dCas9 expression. 0.03 µg/ml dalbavancin was added where indicated.

## Legends supplementary tables

**Table S1. Sequences and off-target analysis of sgRNAs in the CRISPRi library.** sgRNAs are numbered 1-1928 and the targeted locus tag is indicated along with the full 20-bp sgRNA sequences and the 12 bp core sequences. The full off-target analysis against the NCTC8325-genome (CP000253) is also shown. See de Bakker et al (23) and https://github.com/veeninglab/CRISPRi-seq for full explanation of column headings.

**Table S2. sgRNA targets in the NCTC8325 genome**. A list of all locus tags in the NCTC8325 genome along with the corresponding sgRNA targeting each of them. The transcriptional unit (as defined by Mäder et al. (38)), the gene name, position of the start and end of the gene (in the NCTC8325-genome), and which strand the gene is located at (1 or -1) are also shown.

**Table S3. Non-targeted locus tags in the NCTC8325 genome**. A list of the locus tags not targeted by any sgRNA. Position of the start and end of the gene (in the NCTC8325-genome) and which strand the gene is located at (1 or -1) are shown. The reason for the lack of targeting sgRNA is also indicated.

**Table S4. sgRNA targets in *S. aureus* MRSA252**. All sgRNAs in the library and their targets in *S. aureus* MRSA252. See de Bakker et al (23) for full explanation of column headings.

**Table S5. sgRNA targets in *S. aureus* RF122**. All sgRNAs in the library and their targets in *S. aureus* RF122. See de Bakker et al (23) for full explanation of column headings.

**Table S6. sgRNA targets in *S. aureus* Mu50**. All sgRNAs in the library and their targets in *S. aureus* Mu50. See de Bakker et al (23) for full explanation of column headings.

**Table S7. sgRNA targets in *S. aureus* Newman**. All sgRNAs in the library and their targets in *S. aureus* Newman. See de Bakker et al (23) for full explanation of column headings.

**Table S8. sgRNA targets in *S. aureus* JE2**. All sgRNAs in the library and their targets in *S. aureus* JE2. See de Bakker et al (23) for full explanation of column headings.

**Table S9. sgRNA targets in *S. aureus* COL**. All sgRNAs in the library and their targets in *S. aureus* COL. See de Bakker et al (23) for full explanation of column headings.

**Table S10. sgRNA targets in *S. aureus* DSM2031**. All sgRNAs in the library and their targets in *S. aureus* DSM2031. See de Bakker et al (23) for full explanation of column headings.

**Table S11. Raw counts NCTC8325-4 CRISPRi-seq.**

**Table S12. Essentialome of NCTC8325-4 as determined by CRISPRi-seq**. Results from the 12-generation experiment and 20-generation experiment are shown, including the log2 fold change (log2FC) between the induced and non-induced condition, the adjusted p-value (padj) and conclusion about essentiality. Whether each sgRNA is considered conserved essential based on previous transposon mutagenesis screens is also is also indicated.

**Table S13. New essentials in NCTC8325-4**. sgRNAs essential in the CRISPRi-seq screens, but not be reported as essential in previous transposon mutagenesis screens. The sgRNAs and their targeted genes are indicated. The possibility for the sgRNAs to affect neighbouring essential genes is also indicated.

**Table S14. Raw counts Newman CRISPRi-seq.**

**Table S15.** Essentialome of *S. aureus* Newman by CRISPRi-seq. Results from the 12-generation CRISPRi-seq experiment in *S. aureus* Newman (shaded in orange), including the log_2_ fold change (log2FC) between the induced and non-induced condition, the adjusted p-value (padj) and conclusion about essentiality. The results of the corresponding 12-generation experiment in NCTC832-4 are also included for comparison (as in Table S12).

**Table S16. Raw counts NCTC8325-4 dalbavancin CRISPRi-seq.**

**Table S17. CRISPRi-seq screen to identify genes influencing dalbavancin susceptibility**. Log_2_ fold-changes between treated and non-treated samples are shown for each sgRNA as well as the corresponding p-values (padj). Essential; genes where knockdown is expected to give increased sensitivity, costly; knockdown expected to give reduced sensitivity.

**Table S18. List of strains used in this study.**

**Table S19. List of primers used in this study.**

**Table S20. sgRNA oligos used in the library**.

## References

1. Murray CJL, Ikuta KS, Sharara F, Swetschinski L, Robles Aguilar G, Gray A, Han C, Bisignano C, Rao P, Wool E, Johnson SC, Browne AJ, Chipeta MG, Fell F, Hackett S, Haines-Woodhouse G, Kashef Hamadani BH, Kumaran EAP, McManigal B, Agarwal R, Akech S, Albertson S, Amuasi J, Andrews J, Aravkin A, Ashley E, Bailey F, Baker S, Basnyat B, Bekker A, Bender R, Bethou A, Bielicki J, Boonkasidecha S, Bukosia J, Carvalheiro C, Castañeda-Orjuela C, Chansamouth V, Chaurasia S, Chiurchiù S, Chowdhury F, Cook AJ, Cooper B, Cressey TR, Criollo-Mora E, Cunningham M, Darboe S, Day NPJ, De Luca M, Dokova K, et al. 2022. Global burden of bacterial antimicrobial resistance in 2019: a systematic analysis. The Lancet 399:629–655.

2. Butler MS, Hansford KA, Blaskovich MAT, Halai R, Cooper MA. 2014. Glycopeptide antibiotics: Back to the future. J Antibiot 67:631–644.

3. Stefani S, Campanile F, Santagati M, Mezzatesta ML, Cafiso V, Pacini G. 2015. Insights and clinical perspectives of daptomycin resistance in *Staphylococcus aureus*: A review of the available evidence. Int J Antimicrob Agents 46:278–289.

4. van Groesen E, Innocenti P, Martin NI. 2022. Recent advances in the development of semisynthetic glycopeptide antibiotics: 2014–2022. ACS Infectious Diseases 8:1381–1407.

5. Coe KA, Lee W, Stone MC, Komazin-Meredith G, Meredith TC, Grad YH, Walker S. 2019. Multi-strain Tn-Seq reveals common daptomycin resistance determinants in *Staphylococcus aureus*. PLoS Pathog 15:e1007862.

6. Qi LS, Larson MH, Gilbert LA, Doudna JA, Weissman JS, Arkin AP, Lim WA. 2013. Repurposing CRISPR as an RNA-guided platform for sequence-specific control of gene expression. Cell 152:1173–83.

7. Bikard D, Jiang W, Samai P, Hochschild A, Zhang F, Marraffini LA. 2013. Programmable repression and activation of bacterial gene expression using an engineered CRISPR-Cas system. Nucleic Acids Res 41:7429–37.

8. Myrbråten IS, Stamsås GA, Chan H, Morales Angeles D, Knutsen TM, Salehian Z, Shapaval V, Straume D, Kjos M. 2022. SmdA is a novel cell morphology determinant in *Staphylococcus aureus*. mBio 13:e0340421.

9. Reed P, Sorg M, Alwardt D, Serra L, Veiga H, Schäper S, Pinho MG. 2022. A CRISPRi-based genetic resource to study essential *Staphylococcus aureus* genes. bioRxiv doi:10.1101/2022.10.31.514627:2022.10.31.514627.

10. Dong X, Jin Y, Ming D, Li B, Dong H, Wang L, Wang T, Wang D. 2017. CRISPR/dCas9-mediated inhibition of gene expression in *Staphylococcus aureus*. J Microbiol Methods 139:79–86.

11. Zhao C, Shu X, Sun B. 2017. Construction of a gene knockdown system based on catalytically inactive (“dead”) Cas9 (dCas9) in *Staphylococcus aureus*. Appl Environ Microbiol 83:e00291–17.

12. Stamsås GA, Myrbråten I, Straume D, Salehian Z, Veening J-W, Håvarstein LS, Kjos M. 2018. CozEa and CozEb play overlapping and essential roles in controlling cell division in *Staphylococcus aureus*. Mol Microbiol 109:615–632.

13. Cui L, Vigouroux A, Rousset F, Varet H, Khanna V, Bikard D. 2018. A CRISPRi screen in *E. coli* reveals sequence-specific toxicity of dCas9. Nat Commun 9:1912.

14. Rousset F, Cui L, Siouve E, Becavin C, Depardieu F, Bikard D. 2018. Genome-wide CRISPR-dCas9 screens in *E. coli* identify essential genes and phage host factors. PLoS Genet 14:e1007749.

15. Wang T, Guan C, Guo J, Liu B, Wu Y, Xie Z, Zhang C, Xing X-H. 2018. Pooled CRISPR interference screening enables genome-scale functional genomics study in bacteria with superior performance. Nat Comm 9:2475.

16. Liu X, Kimmey JM, Matarazzo L, de Bakker V, Van Maele L, Sirard JC, Nizet V, Veening JW. 2021. Exploration of bacterial bottlenecks and *Streptococcus pneumoniae* pathogenesis by CRISPRi-seq. Cell Host Microbe 29:107–120 e6.

17. Knoops A, Waegemans A, Lamontagne M, Decat B, Mignolet J, Veening JW, Hols P. 2022. A Genome-Wide CRISPR Interference Screen Reveals an StkP-Mediated Connection between Cell Wall Integrity and Competence in Streptococcus salivarius. mSystems 0:e00735–22.

18. Hawkins JS, Silvis MR, Koo BM, Peters JM, Osadnik H, Jost M, Hearne CC, Weissman JS, Todor H, Gross CA. 2020. Mismatch-CRISPRi reveals the co-varying expression-fitness relationships of essential genes in *Escherichia coli* and *Bacillus subtilis*. Cell Syst 11:523– 535.e9.

19. de Wet TJ, Gobe I, Mhlanga MM, Warner DF. 2018. CRISPRi-Seq for the identification and characterisation of essential mycobacterial genes and transcriptional units. bioRxiv doi:10.1101/358275:358275.

20. Ward RD, Tran JS, Banta AB, Bacon EE, Rose WE, Peters JM. 2023. Essential gene knockdowns reveal genetic vulnerabilities and antibiotic sensitivities in *Acinetobacter baumannii*. bioRxiv doi:10.1101/2023.08.02.551708.

21. Lee HH, Ostrov N, Wong BG, Gold MA, Khalil AS, Church GM. 2019. Functional genomics of the rapidly replicating bacterium *Vibrio natriegens* by CRISPRi. Nat Microbiol 4:1105–1113.

22. Jiang W, Oikonomou P, Tavazoie S. 2020. Comprehensive genome-wide perturbations via CRISPR adaptation reveal complex genetics of antibiotic sensitivity. Cell 180:1002–1017.e31.

23. de Bakker V, Liu X, Bravo AM, Veening JW. 2022. CRISPRi-seq for genome-wide fitness quantification in bacteria. Nat Protoc 17:252–281.

24. Dewachter L, Dénéréaz J, Liu X, de Bakker V, Costa C, Baldry M, Sirard J-C, Veening J-W. 2022. Amoxicillin-resistant *Streptococcus pneumoniae* can be resensitized by targeting the mevalonate pathway as indicated by sCRilecs-seq. eLife 11:e75607.

25. Abbas M, Paul M, Huttner A. 2017. New and improved? A review of novel antibiotics for Gram-positive bacteria. Clinical Microbiology and Infection 23:697–703.

26. Zeng D, Debabov D, Hartsell TL, Cano RJ, Adams S, Schuyler JA, McMillan R, Pace JL. 2016. Approved glycopeptide antibacterial drugs: mechanism of action and resistance. Cold Spring Harbor Perspectives in Medicine 6:a026989.

27. Economou NJ, Nahoum V, Weeks SD, Grasty KC, Zentner IJ, Townsend TM, Bhuiya MW, Cocklin S, Loll PJ. 2012. A carrier protein strategy yields the structure of dalbavancin. J Am Chem Soc 134:4637–4645.

28. Sader HS, Mendes RE, Duncan LR, Pfaller MA, Flamm RK. 2018. Antimicrobial activity of dalbavancin against *Staphylococcus aureus* with decreased susceptibility to glycopeptides, daptomycin, and/or linezolid from U.S. Medical Centers. Antimicrob Agents Chemother 62.

29. Andes D, Craig William A. 2007. *In vivo* pharmacodynamic activity of the glycopeptide Dalbavancin. Antimicrob Agents Chemother 51:1633–1642.

30. Morrisette T, Miller MA, Montague BT, Barber GR, McQueen RB, Krsak M. 2019. On- and off-label utilization of dalbavancin and oritavancin for Gram-positive infections. J Antimicrob Chemother 74:2405–2416.

31. Boucher HW, Wilcox M, Talbot GH, Puttagunta S, Das AF, Dunne MW. 2014. Once-weekly dalbavancin versus daily conventional therapy for skin infection. N Engl J Med 370:2169–2179.

32. Raad I, Darouiche R, Vazquez J, Lentnek A, Hachem R, Hanna H, Goldstein B, Henkel T, Seltzer E. 2005. Efficacy and safety of weekly dalbavancin therapy for catheter-related bloodstream infection caused by Gram-positive pathogens. Clin Infect Dis 40:374–380.

33. Werth BJ, Jain R, Hahn A, Cummings L, Weaver T, Waalkes A, Sengupta D, Salipante SJ, Rakita RM, Butler-Wu SM. 2018. Emergence of dalbavancin non-susceptible, vancomycin-intermediate *Staphylococcus aureus* (VISA) after treatment of MRSA central line-associated bloodstream infection with a dalbavancin- and vancomycin-containing regimen. Clin Microbiol Infect 24:429.e1–429.e5.

34. Gardete S, Kim C, Hartmann BM, Mwangi M, Roux CM, Dunman PM, Chambers HF, Tomasz A. 2012. Genetic pathway in acquisition and loss of vancomycin resistance in a methicillin resistant *Staphylococcus aureus* (MRSA) strain of clonal type USA300. PLoS Pathog 8:e1002505.

35. Howden BP, Davies J, K., Johnson P, D. R., Stinear T, P., Grayson ML. 2010. Reduced vancomycin susceptibility in *Staphylococcus aureus*, including vancomycin-intermediate and heterogeneous vancomycin-intermediate strains: resistance mechanisms, laboratory detection, and clinical implications. Clin Microbiol Rev 23:99–139.

36. Werth BJ, Ashford NK, Penewit K, Waalkes A, Holmes EA, Ross DH, Shen T, Hines KM, Salipante SJ, Xu L. 2021. Dalbavancin exposure in vitro selects for dalbavancin-non-susceptible and vancomycin-intermediate strains of methicillin-resistant *Staphylococcus aureus*. Clin Microbiol Infect 27:910.e1–910.e8.

37. Liew AT, Theis T, Jensen SO, Garcia-Lara J, Foster SJ, Firth N, Lewis PJ, Harry EJ. 2011. A simple plasmid-based system that allows rapid generation of tightly controlled gene expression in *Staphylococcus aureus*. Microbiology 157:666–76.

38. Mader U, Nicolas P, Depke M, Pane-Farre J, Debarbouille M, van der Kooi-Pol MM, Guerin C, Derozier S, Hiron A, Jarmer H, Leduc A, Michalik S, Reilman E, Schaffer M, Schmidt F, Bessieres P, Noirot P, Hecker M, Msadek T, Volker U, van Dijl JM. 2016. *Staphylococcus aureus* transcriptome architecture: from laboratory to infection-mimicking conditions. PLoS Genet 12:e1005962.

39. Kanehisa M, Goto S, Kawashima S, Nakaya A. 2002. The KEGG databases at GenomeNet. Nucleic Acids Res 30:42–6.

40. Santiago M, Matano LM, Moussa SH, Gilmore MS, Walker S, Meredith TC. 2015. A new platform for ultra-high density *Staphylococcus aureus* transposon libraries. BMC Genomics 16:252.

41. Valentino MD, Foulston L, Sadaka A, Kos VN, Villet RA, Santa Maria J, Jr., Lazinski DW, Camilli A, Walker S, Hooper DC, Gilmore MS. 2014. Genes contributing to *Staphylococcus aureus* fitness in abscess- and infection-related ecologies. MBio 5:e01729–14.

42. Chaudhuri RR, Allen AG, Owen PJ, Shalom G, Stone K, Harrison M, Burgis TA, Lockyer M, Garcia-Lara J, Foster SJ, Pleasance SJ, Peters SE, Maskell DJ, Charles IG. 2009. Comprehensive identification of essential *Staphylococcus aureus* genes using Transposon-Mediated Differential Hybridisation (TMDH). BMC Genomics 10:291.

43. Lopez S, Hackbarth C, Romanò G, Trias J, Jabes D, Goldstein BP. 2005. In vitro antistaphylococcal activity of dalbavancin, a novel glycopeptide. J Antimicrob Chemother 55:ii21-ii24.

44. Blake KL, O’Neill AJ. 2013. Transposon library screening for identification of genetic loci participating in intrinsic susceptibility and acquired resistance to antistaphylococcal agents. J Antimicrob Chemother 68:12–6.

45. Downer R, Roche F, Park PW, Mecham RP, Foster TJ. 2002. The Elastin-binding Protein of Staphylococcus aureus(EbpS) Is Expressed at the Cell Surface as an Integral Membrane Protein and Not as a Cell Wall-associated Protein*. Journal of Biological Chemistry 277:243–250.

46. Downer R, Roche F, Park PW, Mecham RP, Foster TJ. 2002. The elastin-binding protein of *Staphylococcus aureus* (EbpS) Is expressed at the cell surface as an inegral membrane protein and not as a cell wall-associated protein. J Biol Chem 277:243–250.

47. Park PW, Rosenbloom J, Abrams WR, Rosenbloom J, Mecham RP. 1996. Molecular Cloning and Expression of the Gene for Elastin-binding Protein (ebpS) in Staphylococcus aureus*. Journal of Biological Chemistry 271:15803–15809.

48. Nakakido M, Aikawa C, Nakagawa I, Tsumoto K. 2014. The staphylococcal elastin-binding protein regulates zinc-dependent growth/biofilm formation. J Biochem 156:155–162.

49. Chan YG, Frankel MB, Missiakas D, Schneewind O. 2016. SagB glucosaminidase Is a determinant of *Staphylococcus aureus* glycan chain length, antibiotic susceptibility, and protein secretion. J Bacteriol 198:1123–36.

50. Schaefer K, Owens TW, Page JE, Santiago M, Kahne D, Walker S. 2021. Structure and reconstitution of a hydrolase complex that may release peptidoglycan from the membrane after polymerization. Nat Microbiol 6:34–43.

51. DeMars Z, Bose JL. 2018. Redirection of metabolism in response to fatty acid kinase in *Staphylococcus aureus*. J Bacteriol 200:e00345–18.

52. Sieradzki K, Tomasz A. 2006. Inhibition of the autolytic system by vancomycin causes mimicry of vancomycin-intermediate *Staphylococcus aureu*s-type resistance, cell concentration dependence of the MIC, and antibiotic tolerance in vancomycin-susceptible *S. aureus*. Antimicrob Agents Chemother 50:527–533.

53. Finan JE, Archer GL, Pucci MJ, Climo MW. 2001. Role of penicillin-binding protein 4 in expression of vancomycin resistance among clinical isolates of oxacillin-resistant *Staphylococcus aureus*. Antimicrob Agents Chemother 45:3070–5.

54. Sieradzki K, Pinho MG, Tomasz A. 1999. Inactivated *pbp4* in highly glycopeptide-resistant laboratory mutants of *Staphylococcus aureus*. J Biol Chem 274:18942–6.

55. Chatterjee SS, Chen L, Gilbert A, da Costa TM, Nair V, Datta SK, Kreiswirth BN, Chambers HF. 2017. PBP4 mediates β-Lactam resistance by altered function. Antimicrob Agents Chemother 61:e00932–17.

56. Masters EA, de Mesy Bentley KL, Gill AL, Hao SP, Galloway CA, Salminen AT, Guy DR, McGrath JL, Awad HA, Gill SR, Schwarz EM. 2020. Identification of penicillin binding protein 4 (PBP4) as a critical factor for *Staphylococcus aureus* bone invasion during osteomyelitis in mice. PLoS Pathog 16:e1008988.

57. Abdul-Mutakabbir JC, Kebriaei R, Stamper KC, Sheikh Z, Maassen PT, Lev KL, Rybak MJ. 2020. Dalbavancin, vancomycin and daptomycin alone and in combination with cefazolin against resistant phenotypes of *Staphylococcus aureus* in a pharmacokinetic/pharmacodynamic model. Antibiotics (Basel) 9:696.

58. Kebriaei R, Rice SA, Singh NB, Stamper KC, Nguyen L, Sheikh Z, Rybak MJ. 2020. Combinations of (lipo)glycopeptides with beta-lactams against MRSA: susceptibility insights. J Antimicrob Chemother 75:2894–2901.

59. Kebriaei R, Rice SA, Stamper KC, Rybak MJ. 2019. Dalbavancin alone and in combination with ceftaroline against four different phenotypes of *Staphylococcus aureus* in a simulated pharmacodynamic/pharmacokinetic model. Antimicrob Agents Chemother 63:e01743–18.

60. Xhemali X, Smith JR, Kebriaei R, Rice SA, Stamper KC, Compton M, Singh NB, Jahanbakhsh S, Rybak MJ. 2019. Evaluation of dalbavancin alone and in combination with β-lactam antibiotics against resistant phenotypes of *Staphylococcus aureus*. J Antimicrob Chemother 74:82–86.

61. Parsons JB, Broussard TC, Bose JL, Rosch JW, Jackson P, Subramanian C, Rock CO. 2014. Identification of a two-component fatty acid kinase responsible for host fatty acid incorporation by *Staphylococcus aureus*. Proc Natl Acad Sci U S A 111:10532–7.

62. Frlan R. 2022. An evolutionary conservation and druggability analysis of enzymes belonging to the bacterial Shikimate pathway. Antibiotics (Basel) 11:675.

63. Lofblom J, Kronqvist N, Uhlen M, Stahl S, Wernerus H. 2007. Optimization of electroporation-mediated transformation: *Staphylococcus carnosus* as model organism. J Appl Microbiol 102:736–747.

64. Sorg RA, Gallay C, Van Maele L, Sirard JC, Veening JW. 2020. Synthetic gene-regulatory networks in the opportunistic human pathogen *Streptococcus pneumoniae*. Proc Natl Acad Sci U S A 117:27608–27619.

65. Arnaud M, Chastanet A, Debarbouille M. 2004. New vector for efficient allelic replacement in naturally nontransformable, low-GC-content, Gram-positive bacteria. Appl Environ Microbiol 70:6887–6891.

66. Porcellato D, Skeie SB. 2016. Bacterial dynamics and functional analysis of microbial metagenomes during ripening of Dutch-type cheese. Int Dairy J 61:182–188.

67. Zhang J, Kobert K, Flouri T, Stamatakis A. 2013. PEAR: a fast and accurate Illumina Paired-End reAd mergeR. Bioinformatics 30:614–620.

68. Bravo AM, Typas A, Veening JW. 2022. 2FAST2Q: a general-purpose sequence search and counting program for FASTQ files. PeerJ 10:e14041.

69. Love MI, Huber W, Anders S. 2014. Moderated estimation of fold change and dispersion for RNA-seq data with DESeq2. Genome Biol 15:550.

70. Reichmann NT, Pinho MG. 2017. Role of SCCmec type in resistance to the synergistic activity of oxacillin and cefoxitin in MRSA. Sci Rep 7:6154.

