## Supplemental Tables S18-S19 for "Genome-wide CRISPRi screens reveal the essentialome and determinants for susceptibility to dalbavancin in *Staphylococcus aureus*"

**Table S18. List of *S. aureus* strains in this study**

| **Strain name** | **Genotype** | **Reference** |
| --- | --- | --- |
| *E. coli* |  |  |
| IM08B | DH10B, Δ*dcm*, Phelp-*hsdMS*, PN25-*hsdS* (strain expressing the S. aureus CC8 specific methylation genes) | (1) |
| VL2336 | Read1-P3-BsmBI-mCherry-BsmBI-dCas9 handle-Sp_ter_-cat, Amp^R^ | This work |
| *S. aureus* |  |  |
| NCTC8325-4 | Derivative of NCTC8325, cured of prophages | (2) |
| Newman | Clinical isolate (ATCC 25904), rsbU+ | (3) |
| MK1465 | NCTC8325-4, pLOW-dCas9, Ery^R^ | (4) |
| MH225 | NCTC8325-4, pLOW-P_spac2_-dCas9, Ery^R^ | This work |
| MH226 | Newman, pLOW2-P_spac2_-dCas9, Ery^R^ | This work |
| MK1482 | NCTC8325-4, GFP^+^ | This work |
| MH220 | MK1482, pLOW-P_spac2_-dCas9, pVL2336-sgRNA(gfp), Ery^R^, Cam^R^. | This work |
| MH221 | MK1482, pLOW-dCas9, pVL2336-sgRNA(gfp), Ery^R^, Cam^R^. | This work |
| MM75 | MH225, pVL2336-sgRNA(notarget). Ery^R^, Cam^R^. | This work |
| MH270 | MH225, pVL2336-sgRNA(SAOUHSC_00685) | This work |
| MH264 | MH225, pVL2336-sgRNA(SAOUHSC_00567) | This work |
| MH275 | MH225, pVL2336-sgRNA(dltA). Ery^R^, Cam^R^. | This work |
| AHF59 | MH225, pVL2336-sgRNA(rpsF). Ery^R^, Cam^R^. | This work |
| AHF60 | MH225, pVL2336-sgRNA(vraF). Ery^R^, Cam^R^. | This work |
| AHF56 | MH225, pVL2336-sgRNA(ezrA). Ery^R^, Cam^R^. | This work |
| AHF61 | MH225, pVL2336-sgRNA(nrdF). Ery^R^, Cam^R^. | This work |
| MK1736 | MH225, pVL2336-sgRNA(SAOUHSC_00678) | This work |
| MK1737 | MH225, pVL2336-sgRNA(SAOUHSC_00892) | This work |
| AHF64 | MH225, pVL2336-sgRNA(pbp4). Ery^R^, Cam^R^. | This work |
| AHF54 | MH225, pVL2336-sgRNA(kapB). Ery^R^, Cam^R^. | This work |
| MK1661 | MH225, pVL2336-sgRNA(ebpS). Ery^R^, Cam^R^. | This work |
| MK1662 | MH225, pVL2336-sgRNA(SAOUHSC_00659). Ery^R^, Cam^R^. | This work |
| MK1663 | MH225, pVL2336-sgRNA(mvaS). Ery^R^, Cam^R^. | This work |
| MK1664 | MH225, pVL2336-sgRNA(sgtB). Ery^R^, Cam^R^. | This work |
| MK1667 | MH225, pVL2336-sgRNA(mvaK2). Ery^R^, Cam^R^. | This work |
| MK1654 | MH225, pVL2336-sgRNA(sagB). Ery^R^, Cam^R^. | This work |
| MK1655 | MH225, pVL2336-sgRNA(aroB). Ery^R^, Cam^R^. | This work |
| MK1656 | MH225, pVL2336-sgRNA(aroA). Ery^R^, Cam^R^. | This work |
| MK1657 | MH225, pVL2336-sgRNA(aroC). Ery^R^, Cam^R^. | This work |
| MK1658 | MH225, pVL2336-sgRNA(aroD). Ery^R^, Cam^R^. | This work |
| MK1659 | MH225, pVL2336-sgRNA(aroK). Ery^R^, Cam^R^. | This work |
| MK1660 | MH225, pVL2336-sgRNA(aroA2). Ery^R^, Cam^R^. | This work |
| MK1560 | NCTC8325-4, Δ*pbp4*::spc, Spc^R^ | This work |
| MK1718 | NCTC8325-4, Δ*kapB*::spc. Spc^R^ | This work |

Ery^R^ = erytromycinresistent, Cam^R^ = chloramphenicol resistant, Spc^R^ = spektinimycinresistent

**Table S19. List of oligos**

| **Oligo name** | **Sequence (5’ – 3’)** |
| --- | --- |
| mvh54_Pspac_Rev | AGCAGAGGTTGTTACTGCTC |
| mvh55_intro_lacO_Pspac | GAGCAGTAACAACCTCTGCTAATTGTGAGCGCTCACAATTCTGAAAAATTTTGCAAAAAGTTGT |
| mk411_pbp4_up_F_NcoI | ACGTCCATGGAAATAAGACAACGAGTCATGGA |
| mk412_pbp4_up_R_overaad | GATCCTAGGTGGGCCCAATCCTTGTCGCTGGATTAGCAC |
| mk413_pbp4_down_F_overaad | CTCGAGCGGCCGCATAGTGTAGCTTGTGCATATGGTGTC |
| mk414_pbp4_down_R_BamHI | TCGAGGATCCAACATGATTAGTGACCCAACTA |
| mk188_aad9_up_F | ATTGGGCCCACCTAGGATC |
| mk189_aad9_down_R | ACTATGCGGCCGCTCGAG |
| AHF13_kapB_up_F_bsaI | TTGGCAGGTCTCCCTATGCGAAGCCCAGAATAATGTAAT |
| AHF14_kapB_up_R_bsaI | TTGGCAGGTCTCCGGACGTGTTTCTTCTTATTTAAATACTTCTG |
| AHF15_aad9_F_bsaI | TTGGCAGGTCTCGGTCCCCACCTAGGATCGAATCCC |
| AHF16_aad9_R_bsaI | TTGGCAGGTCTCGGCGAGGCCGCGGTAATAAAC |
| AHF17_kapB_dwn_F_bsaI | TTGGCAGGTCTCCTCGCCTCCCTTAAAAGTATGTTAATATATATGTATCA |
| AHF18_kapB_dwn_R_bsaI | TTGGCAGGTCTCCCTGCTAAAAACATAAAAAGCAACCTCAACTAT |
| AHF19_aad9_F_bsaI | TGTCGAGGTCTCCTGATTGGGCCCACCTAGGATCG |
| AHF19_aad9_R_bsaI | TGTCGAGGTCTCCTACTATGCGGCCGCTCG |
| OVL2127_pCG248-MutBsmBI-F | ATATCGTCTCAGGTTAATGTCATGATAATAATGGTTTCTT |
| OVL2128_pCG248-MutBsmBI-R | ATATCGTCTCGAACCTCACAGCTTGTCTGTAAGCGGAT |
| OVL2152_Read1-IF-F | TTCGGTCGACAGATCTTCGTCGGCA |
| OVL2153_P7-IF-R | CGGTACCCGGGGATCCCAAGCAGAAGACGGCATACGAG |

**References**

1.	Monk IR, Tree JJ, Howden BP, Stinear TP, Foster TJ. 2015. Complete bypass of restriction systems for major *Staphylococcus aureus* lineages. MBio 6:e00308.

2.	Novick R. 1967. Properties of a cryptic high-frequency transducing phage in *Staphylococcus aureus*. Virology 33:155-166.

3.	Duthie ES, Lorenz LL. 1952. Staphylococcal coagulase; mode of action and antigenicity. J Gen Microbiol 6:95-107.

4.	Stamsås GA, Myrbråten I, Straume D, Salehian Z, Veening J-W, Håvarstein LS, Kjos M. 2018. CozEa and CozEb play overlapping and essential roles in controlling cell division in *Staphylococcus aureus*. Mol Microbiol 109:615-632.
